# A spatially constrained independent component analysis jointly informed by structural and functional network connectivity

**DOI:** 10.1101/2023.08.13.553101

**Authors:** Mahshid Fouladivanda, Armin Iraji, Lei Wu, Theodorus G.M. van Erp, Aysenil Belger, Faris Hawamdeh, Godfrey D. Pearlson, Vince D. Calhoun

## Abstract

There are a growing number of neuroimaging studies motivating joint structural and functional brain connectivity. Brain connectivity of different modalities provides insight into brain functional organization by leveraging complementary information, especially for brain disorders such as schizophrenia. In this paper, we propose a multi-modal independent component analysis (ICA) model that utilizes information from both structural and functional brain connectivity guided by spatial maps to estimate intrinsic connectivity networks (ICNs). Structural connectivity is estimated through whole-brain tractography on diffusion-weighted MRI (dMRI), while functional connectivity is derived from resting-state functional MRI (rs-fMRI). The proposed structural-functional connectivity and spatially constrained ICA (sfCICA) model estimates ICNs at the subject level using a multi-objective optimization framework. We evaluated our model using synthetic and real datasets (including dMRI and rs-fMRI from 149 schizophrenia patients and 162 controls). Multi-modal ICNs revealed enhanced functional coupling between ICNs with higher structural connectivity, improved modularity, and network distinction, particularly in schizophrenia. Statistical analysis of group differences showed more significant differences in the proposed model compared to the unimodal model. In summary, the sfCICA model showed benefits from being jointly informed by structural and functional connectivity. These findings suggest advantages in simultaneously learning effectively and enhancing connectivity estimates using structural connectivity.

## 1. Introduction

The human brain can be modeled as a distinct functional unit that exhibits a temporally synchronized pattern known as intrinsic connectivity networks (ICNs) (Genon et al., 2018; Iraji et al., 2020). These ICNs interact with each other, collectively representing brain function. Consequently, accurate estimation of ICNs is a critical step in studying brain networks because it minimizes the impact of inaccuracies in functional activities (Iraji et al., 2020). One approach for identifying ICNs and gaining insights into brain networks, including brain disorders like schizophrenia (Iraji et al., 2019; Iraji, Faghiri, Fu, Kochunov, et al., 2022; Iraji, Faghiri, Fu, Rachakonda, et al., 2022), is data-driven independent component analysis (ICA) (Calhoun & Adalı, 2012; Calhoun & de Lacy, 2017). This method assumes spatial independence for the temporally synchronized functional units and recovers a set of maximally independent sources from a mixture of unknown source signals without any prior information (Calhoun et al., 2001), using single modality neuroimaging, particularly resting-state functional magnetic resonance imaging (rs-fMRI). Rs-fMRI images dynamically measure the hemodynamic response associated with neural activity in the brain during rest.

ICA has become widely used to partition the brain into spatially overlapping and distinct ICNs at the group level. Group ICNs are estimated using fMRI images from all subjects, then the corresponding subject-level ICNs are obtained. The most common approach is to apply ICA to the data, then use back reconstruction to estimate subject-specific maps (Allen et al., 2011; Calhoun et al., 2001; Erhardt et al., 2011). These approaches are fully data-driven; however, more recently, spatially constrained ICA approaches (Du & Fan, 2013; Lin et al., 2010) have been proposed to estimate ICNs using a set of templates for ICNs as priors. They can be used to build a fully automated approach (e.g., the NeuroMark pipeline (Du et al., 2020)) that does not require post hoc ICN selection or network matching (Du et al., 2020), and also automatically provides ICN ordering, providing modular functional network connectivity. These approaches have been used in many prior studies and offer a fully automated framework that can be integrated within a larger framework (e.g., a containerized version is available from http://trendscenter.org/software/gift, as well as a BIDSapp (Kim et al., 2023)), making them more easily comparable across studies.

Current constrained ICA approaches are unimodal and do not leverage information from other modalities, particularly dMRI images. Although MRI images, such as T1 and T2, include structural information about the brain, dMRI images provide information about the physical paths in the human brain, captured as structural connectivity which drives functional activities of different units. In this study, we investigate a model that estimates functional brain patterns (ICNs) informed by multi-modal data to provide a more complete view of the connectome by employing both structural and functional connectivity. This has many advantages, for example, a connectivity domain ICA approach (Iraji et al., 2016) called joint connectivity matrix ICA has been used to jointly parcellate structural and functional connectivity data, yielding a data-driven structure/function parcellation (Wu & Calhoun, 2023).

One longstanding aspiration in brain research is to link brain function to its underlying architecture, making the interplay between structural connectivity and functional network connectivity (FNC) a fundamental in network neuroscience (Park & Friston, 2013). There have been studies, such as (Calhoun & Sui, 2016) using computational models on the connectivity domain to estimate the human brain functional activities propounding from different modalities, mainly single modality information. Consequently, a fundamental question arises regarding how to model human brain functional activity changes with a deeper understanding of structural and functional information (Batista-García-Ramó & Fernández-Verdecia, 2018; Skudlarski et al., 2010). Several studies have been proposed to fill this gap ((Batista-García-Ramó & Fernández-Verdecia, 2018; Suárez et al., 2020)) by developing different models predicting FNC from structural connectivity using statistical models (Messé et al., 2014; Mišić et al., 2016; Suárez et al., 2020), or communication models based on structural-functional network connectivity (Goñi et al., 2014; Zamani Esfahlani et al., 2022).

In addition, jointly analyzing models is a growing methodology used to investigate the human brain and how structural and functional relate to each other (Calhoun & Sui, 2016; Puxeddu et al., 2022; Zhu et al., 2021). One may provide variations in brain functions at different regions by multilayer network models (Puxeddu et al., 2020; Shine et al., 2016). Multilayer network models have emerged to analyze the human brain employing the concepts of graph theory with multiple viewings of the human brain (Puxeddu et al., 2020; Puxeddu et al., 2022). In parallel, other studies have investigated brain network modules, uncovering intrinsic brain network activations using data-driven models (Calhoun & Adalı, 2012; Yeo et al., 2011), and community detection (Power et al., 2011). Several studies have attempted to develop multi-modal fusion models (Calhoun & Sui, 2016; Sui et al., 2014) incorporating complementary information across modalities into the analysis of ICNs (Batista-García-Ramó & Fernández-Verdecia, 2018; Suárez et al., 2020). Multi-modal fusion can provide additional insights into brain structure and function that are impacted by psychopathology, including identifying which structural or functional aspects of pathology might be linked to human behavior or cognition (Sui et al., 2014), particularly in disorders related to both brain function and structures such as schizophrenia (Calhoun, 2018).

Moreover, information from different modalities can be jointly analyzed to investigate ICNs using ICA. This has been done mostly by focusing on linking group-level spatial features, including joint ICA (jICA) (Calhoun et al., 2006), parallel ICA (pICA) (Liu et al., 2009), parallel ICA with reference (pICAR) (Chen et al., 2013), multi-modal canonical correlation analysis (mCCA)+jICA (Sui et al., 2011), multisite canonical correlation analysis with reference (mCCAR)+jICA (Qi et al., 2018; Shile et al., 2016), and linked ICA (Groves et al., 2011), or jointly optimizing for fMRI networks and covarying networks from structural MRI as in parallel group ICA+ICA.

However, these approaches have not directly integrated structural and functional network connectivity information (Qi et al., 2022; S. L. Qi et al., 2019). To our knowledge, fewer prior works have used structural connectivity to estimate ICNs, as in (Wu & Calhoun, 2023), which performed joint structure/functional parcellation. Specifically, there is a lack of methodologies to guide adaptive ICN estimation by incorporating both structural and functional network connectivity. A common practice in neuroscience is to compress Network connectivity information into nodes and edges to understand functional interactions in the brain (Babaeeghazvini et al., 2021). Thus, using network connectivity helps to drive the model to learn from both structural connectivity (real connections among different functional networks) and FNC (their functional interactions).

In this paper, we introduce a novel adaptive ICA model designed to directly investigate the ICNs. This is achieved by imposing constraints using two different modalities at the subject level. Our approach involves constructing a new multi-modal ICA model that jointly embeds structural and functional network connectivity. This model is supported by previous studies that indicate that structural connectivity forms the foundation of functional connectivity (Honey et al., 2009; Litwińczuk et al., 2022; Stam et al., 2016; Zhu et al., 2021). In addition, we incorporated prior spatial information into our multi-modal ICA model to increase the generalizability of the estimated ICNs. Our model is an iterative pipeline that simultaneously learns from both dMRI and fMRI, along with ICN spatial maps, to identify joint structural-functional independent components.

While prior work (Wu & Calhoun, 2023) applied a connectivity-based model to perform joint data-driven parcellation of dMRI and fMRI, no approaches have attempted to directly link the full spatiotemporal fMRI data with dMRI data in the context of a connectivity constraint at the subject level within a spatially constrained model, thus providing a fully automated approach. Our proposed structural-functional network connectivity and spatially constrained ICA model (sfCICA) identifies ICNs that are simultaneously optimized to be maximally spatially independent, influenced by both structural and functional connectivity. Thus, the learning procedure to estimate subject-level ICNs is informed by the structural and functional network connectivity information of each subject. We first validated the approach employing synthetic data and then applied the proposed model to a real dataset, including controls and patients with schizophrenia. The results demonstrate that the proposed method can effectively impose prior information on ICNs while jointly learning from structural-functional network connectivity in both synthetic and real human brain data. Furthermore, the identified ICNs exhibit an increased ability to distinguish between healthy controls and individuals with schizophrenia.

## 2. Material and Methods

### 2.1. Structural-functional network connectivity and spatially constrained ICA (sfCICA)

#### Model definition

A multi-modal ICA (sfCICA) model, called *structural and functional network connectivity and spatially constrained ICA* was introduced. Briefly, the sfCICA framework estimates subject-level ICNs by imposing structural network connectivity weights on time course distances (functional network connectivity). The main advantage of the constraints in the sfCICA framework is driving the model to jointly learn from both structural and functional network connectivity at the subject level. Using the sfCICA model, subject-specific ICNs were estimated through multi-modal information, incorporating prior spatial maps for each ICN derived from a standard template.

In summary, we introduced a multi-objective function as described in **Eq. 1**, consisting of three terms. The first and second terms find independent ICNs which are learned from the spatial template. The third term constrains the learning procedure to account for both structural and functional network connectivity.

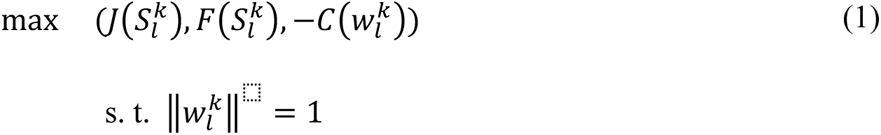

The sfCICA model was proposed in a multi-objective framework, aiming to estimate ICNs by optimizing their independence, through maximizing their non-Gaussianity measured by negentropy or kurtosis 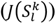 (Hyvarinen, 1999) (**Eq. 2**).

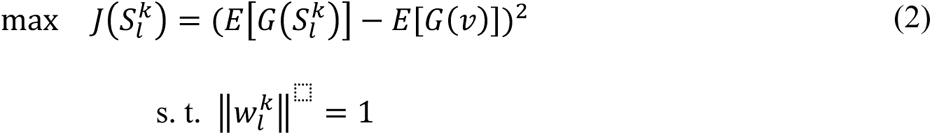

Where, G is any quadratic function, E is the expectation maximization operator, and *v* is a random variable. 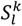 and 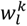 are respectively the spatial map and time course of the *l^th^* ICN (here we use Neuromark_fMRI_1.0 including 53 ICNs) for *k^th^* subject. These are estimated by maximizing the square differences between the expectation maximization of two vectors, as defined in **Eq. 2**. In addition, the similarity of the spatial maps with the prior spatial information was maximized by another cost function 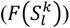 in **Eq. 3**. Thus, independence and similarity to the prior spatial information (here a standard template was used) were jointly maximized during optimization.

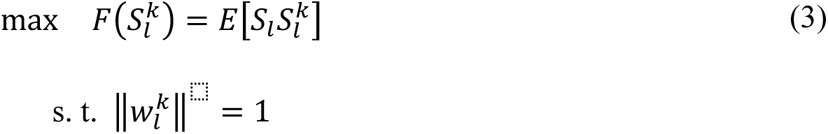

where the *S_l_* denoted the spatial map of the *l^th^* components in the template. Furthermore, we integrate structural connectivity information, measured as the number of streamlines connecting node pairs, into the model as a constraint on the FNC of the estimated time course. To this end, the model is guided by minimizing the weighted distances of the time courses based on structural connectivity. The structural connectivity weights captured from dMRI can reflect existing physical connectivity between functionally related gray matter regions, this means, more fiber bundles between two brain regions (here are ICNs) may result in stronger functional correlations and shorter distances between time courses (Zhu et al., 2021).

Thus, we assumed regions with higher structural connectivity (more fiber tracts) would exhibit closer functional signal activities because of their shorter distance. Prior works (He & Niyogi, 2003; Zhu et al., 2021), led us to reconstruct an objective function 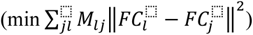) to adjust the functional activity of an ICN by considering other structurally connected ICNs. Here, *M_lj_* represents structural connectivity weights and *FC* denotes the correlation between pairs of ICNs (*l*, *j*). It is important to note that maximizing the FC between ICNs involves minimizing the distance of their time courses. Therefore, we propose the following cost function (**Eq. 4**), in addition to the previous cost functions, to be simultaneously optimized. This is aimed at incorporating information from both structure and function as constraints.

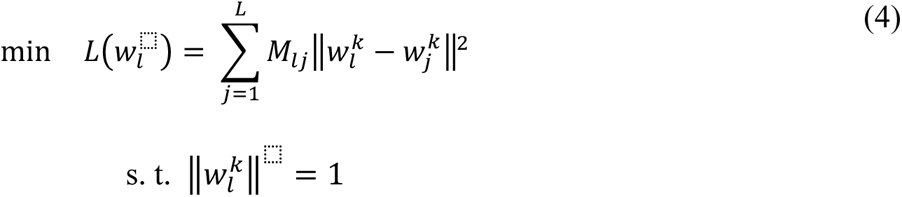

To solve a multi-objective optimization problem, one usually needs to find the Pareto optimal set or its subset and critically evaluate which specific trade-off is more appropriate to the problem under study. In addition, optimizing a linear weighted sum of cost functions is a commonly used method for solving multi-objective optimization problems, which simplifies the problem (Klamroth & Jørgen, 2007). In our study, we utilize a linear weighted approach to address the multi-objective problem described in **Eq. 1**. This allows us to explore various points along the Pareto front by adjusting the weighting values in the linear weighted sum objective function.

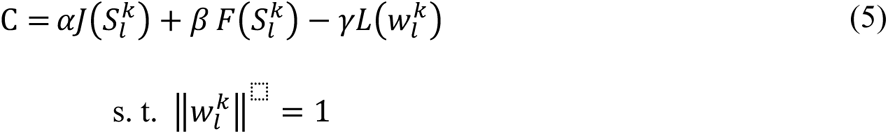

Where, *α*, *β*, and *γ* are constant weight values. To ensure a fair comparison and prevent the optimization from being biased towards a cost function with a larger magnitude, we normalize the cost functions in **Eq. 2-4** as described in (Du & Fan, 2013), and weights (*α*, *β*, and *γ*) are chosen to be equal (0.33). Then a gradient ascent method was used to iteratively converge on an optimal solution. The iterative algorithm for optimizing the cost (C) is derived as follows. Initially, 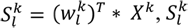, is substituted into **Eq. 6**.

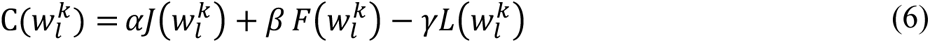

Then, according to the derivatives of each term, we have:

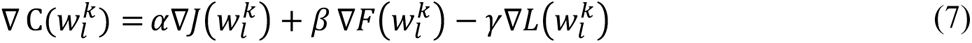

More details regarding the derivative of each term are described in (Du & Fan, 2013). Once the gradient of the objective function of **Eq. 7** is available, we employ the steepest ascent iterative formula:

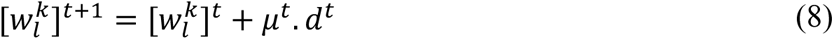

Where 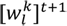 represents the value of 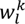 after t+1 iteration, *d^t^* is the normalized 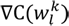 and, *μ^t^* stands for the step-length. To ensure a sufficient increase in the objective function in an inexact line search, *μ^t^* was estimated using the Armijo condition (Jorge Nocedal, 2006).

### 2.2. Evaluation analysis

To evaluate the benefit of leveraging multi-modal information, we compare our model with one of the spatial constrained ICA (CICA) models, known as a multi-objective optimization ICA model (Du et al., 2020), which is a simplified version of the proposed framework when the multi-modal information is not utilized. To this end, first, we used synthetic data to confirm that the model is working in the intended objective defined for the optimization of the time series. Then, we cope to evaluate our hypothesis using real data, capturing better group differences by leveraging multi-modal information.

#### 2.2.1. Synthetic data

##### Simulation

The synthetic dMRI was from FiberCup phantom data (Fillard et al., 2011; Poupon et al., 2008) (https://tractometer.org/fibercup/data/), including 3mm isotropic images in 64 uniformly distributed over a sphere with b-value = 2000 m/s. It also provided 16 brain regions and fiber pathways among the regions to reconstruct structural connectivity, (shown in **Figure 1**). To keep consistency between the simulated dMRI and fMRI data, we used the same sixteen predefined FiberCup regions as a reference to design synthetic fMRI images, similar to (Chu et al., 2018). We simulated synthetic fMRI images using the simTB toolbox (Erhardt et al., 2012), (https://trendscenter.org/software/simtb/).

**Figure 1.**
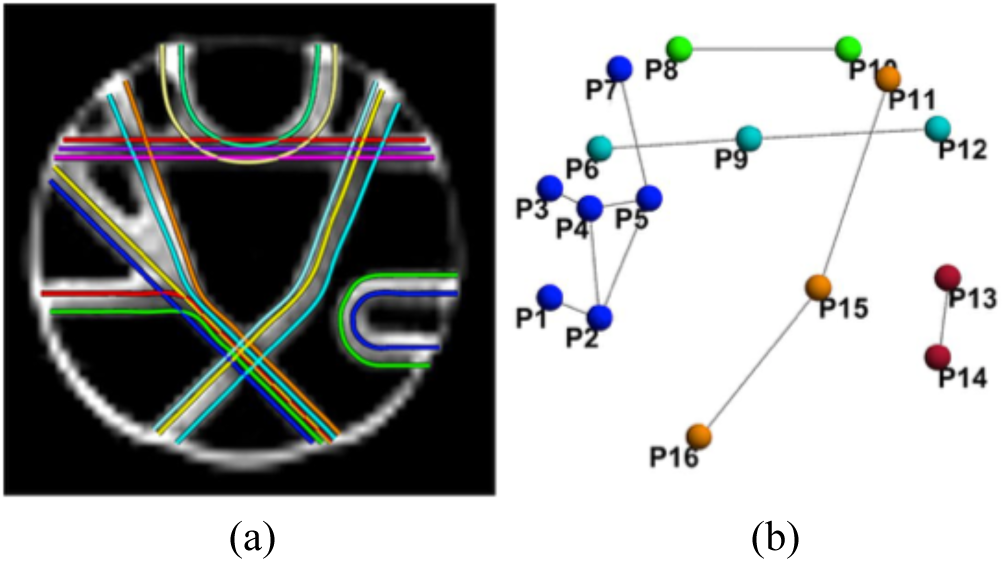
In panel (a) and (b), we present simulated fiber pathways and predefined regions from FiberCup, respectively, for the reconstruction of synthetic structural connectivity.

The simTB toolbox employs a data generation model that assumes spatiotemporal separability, allowing the simulated fMRI data to be represented as the multiplication of spatial maps and time courses. The generated data exhibit realistic dimensions, spatiotemporal activations, and noise characteristics similar to those observed in typical fMRI datasets (Erhardt et al., 2012). Our fMRI simulation consists of 16 ICNs for each of the 30 subjects, evenly divided into 3 distinct groups (10 subjects per group) with varying design parameters (Group A, B and C). Generally, all spatial maps have V = 64 × 64 voxels (where V is the total number of voxels) and time courses are T = 3000 time points in length with a repetition time (TR) of 2 seconds. Spatial maps of the ICNs represented in **Figure 2** correspond to the regions in the FiberCup data shown in **Figure 1**.

**Figure 2.**
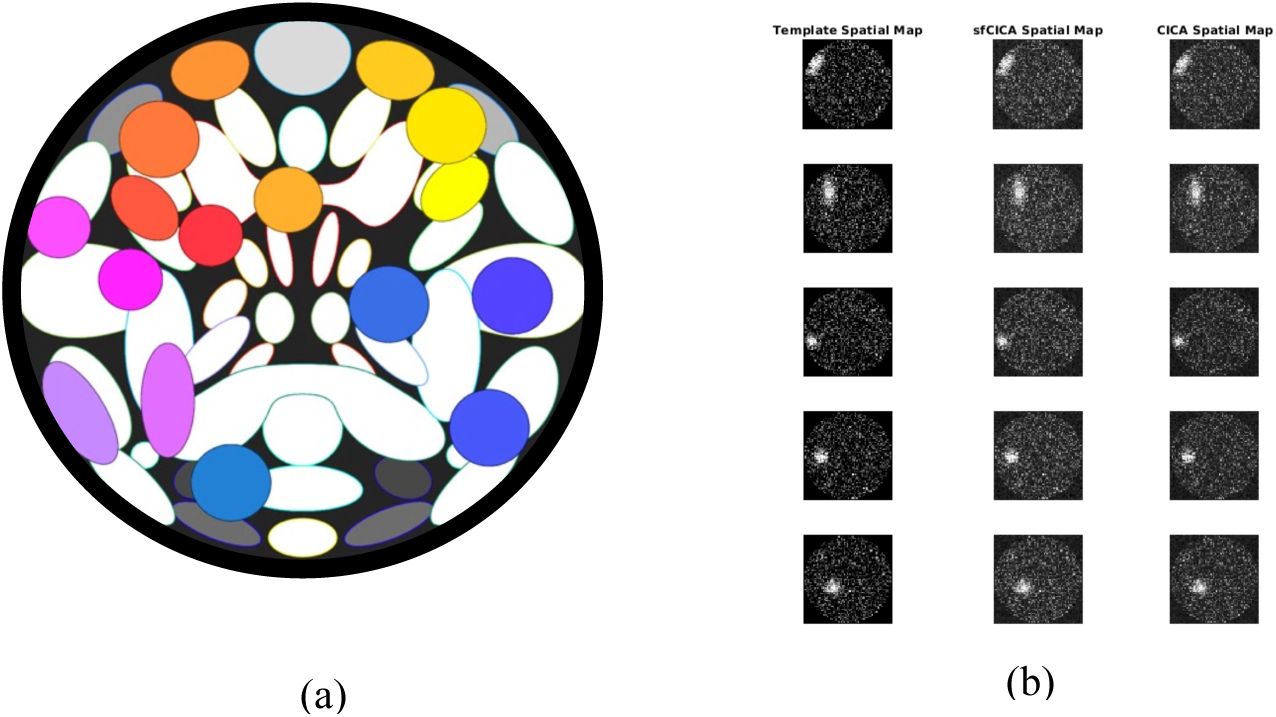
(a) It represents the 16 spatial maps of the ICNs, derived from the FiberCup regions, inserted into the simTB toolbox as a reference to generate synthetic fMRI images. Using the generated synthetic data, we determined an average template, shown in the first column from the left (b), to be used as a prior spatial map. The second and third columns from the left in (b) illustrate examples of estimated ICNs by the sfCICA and CICA model using synthetic fMRI.

Groups A, B, and C differ in four ways, as outlined below. Firstly, the noise level for each group was specifically selected to determine the robustness of the proposed model. For Group A, the mean was 1.7, with a standard deviation of 0.2; for Group B, the mean was 1.5 with a standard deviation of 0.4; and for Group C, the mean was 1.1 with a standard deviation of 0.35. Secondly, to introduce diversity among the ICNs across different Groups, we implemented additional adjustments to both the amplitudes (g_ic_), and shapes of the reference ICNs. For ICN 6 in Group A, we selected a weaker amplitude compared to Group B and C, while Group C had a weaker amplitude for ICNs 8 and 11 compared to Groups A and B. The amplitudes of the ICNs (g_ic_) were modeled on a normal distribution. For instance, the distribution of ICN 6 (g_ic6_) is normal with a mean of 2.5 and a standard deviation of 0.3 for Group A, and a mean of 3.5 with a standard deviation of 0.3 for Group B. Similarly, ICN 8 and 11 in Group C follow a normal distribution with a mean of 2.5 and a standard deviation of 0.3, whereas Groups A and B have a mean of 3.5 and a standard deviation of 0.3. Additionally, each ICN underwent distinct transitions in the x and y directions, as well as rotations, introducing unique changes. Moreover, the ICNs in Groups A and C were expanded by factors of 1.2 and 1.3, respectively, while the ICNs in Group B were contracted by a factor of 0.9.

Then, we constructed a synthetic ICN template to be used as a prior spatial map in the constrained ICA model. The synthetic template was determined by averaging each ICN over all groups and introducing deformation and noise.

##### Evaluation criteria

Our model works on decreasing the weighted distance between time courses of the structurally connected ICNs. To ensure the objective is preserved, we used a distance metric and sparsity to measure the effect of the structural connectivity constraint on TCs. For this purpose, the sum of the distances between an ICN with all other ICs was determined utilizing the Euclidean distance as outlined in **Eq 9.**

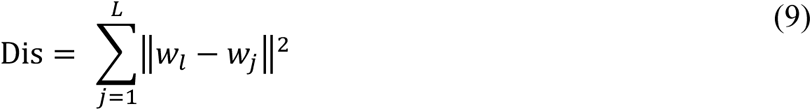

where *w_l_* is the time course of ICN number *l* and *w_j_* is the time courses of all other remaining ICNs. This measure was estimated for both the sfCICA and CICA models.

Sparsity was measured as the number of weak connections to all possible connections in the FNC. To identify weak connections, we first determined an optimal threshold by maximizing the global cost-efficiency of the brain networks (Bassett et al., 2009), (here a threshold of 0.45 was determined). The remaining connections after this threshold were considered strong connections (ST). Then, all FNC networks were thresholded to retain strong connections and sparsity as follows.

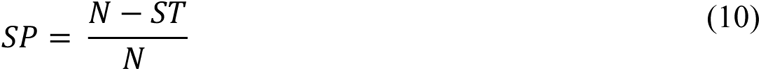

Where ST is the number of strong connections and N is the total number of possible connections. Moreover, the robustness of the model was assessed by varying noise levels across the three distinct simulated groups.

#### 2.2.2. Real data

In this study, the real rs-fMRI and dMRI images were used from the Function Biomedical Informatics Research Network (FBIRN) phase III datasets (Keator et al., 2016). Data collection was at multiple sites of healthy control (HC) subjects or with schizophrenia (SZ) disorder. FBIRN dataset consists of rs-fMRI and dMRI images from 311 age-gender-matched adults (age ranging from 18 to 65 years old). It is 162 healthy controls (HC) including 115 male and 45 female (Avg ± SD age: 37.0 ± 10.9) and 149 schizophrenia (SZ) patients including 115 male and 36 female (Avg ± SD age: 38.7 ± 11.6).

##### Acquisition

Images were scanned at six 3T Siemens Tim Trio System and one 3T GE Discovery MR750 scanner at multiple sites. The acquisition parameters of the rs-fMRI imaging were as follows: FOV of 220 × 220 mm (64 × 64 matrix), TR = 2 s, TE = 30 ms, flip angle (FA) = 77°, 162 volumes, 32 sequential ascending axial slices of 4 mm thickness and 1 mm skip. Subjects had their eyes closed during the resting state scan. For more details see (Keator et al., 2016). For the dMRI images, scanning protocols were settled virtually to ensure equivalent acquisitions with 30 directions of diffusion gradient at b = 800 s/mm^2^ and five measurements of b = 0 (b0) s/mm^2^ (Keator et al., 2016).

##### rs-fMRI preprocessing

The rs-fMRI images were preprocessed via the statistical parametric mapping (SPM12, http://www.fil.ion.ucl.ac.uk/spm/) toolbox in MATLAB 2020b, following the procedure outlined in (Iraji, Fu, et al., 2022). The main steps of the preprocessing included removing the first five fMRI timepoints, motion correction, slice timing correction, image registering to the standard Montreal Neurological Institute (MNI) space, spatial resampling (to 3 × 3 ×3 mm^3^ isotropic voxels), spatial smoothing using Gaussian kernel with a full width at half maximum (FWHM) = 6 mm.

##### dMRI preprocessing

For preprocessing of the dMRI images, first, all images were corrected for eddy current and motion distortions using the eddy package (FSL 6.0 (Jenkinson et al., 2012)) by motion induced signal dropout detection based on b0 volumes and replacement method (Andersson & Sotiropoulos, 2016). Then, a data quality check was performed by an in-house developed algorithm and visual inspection to exclude images with extreme head motion, signal dropout, or noise level (Caprihan et al., 2011; Wu et al., 2015).

An additional step was carried out on preprocessed dMRI images to determine white matter fiber tracts for constructing structural connectivity. First, we used linear regression under the dtifit in the FSL toolbox (Jenkinson et al., 2012) to model a voxel-wise diffusion tensor. The estimated diffusion tensors were employed in deterministic tractography using the CAMINO toolbox (Cook et al., 2006). This procedure involved three stopping criteria: a fixed step size of 0.5mm, an anisotropy threshold of 0.2, and an angular threshold of 60° (Cook et al., 2006). All fiber tracts were extracted in native space. Next, we transformed the native fractional anisotropy (FA) maps into the standard MNI space using advanced normalizations tools (ANTs) (Avants et al., 2009). Then we applied inverted spatial normalization to the Neuromark_fMRI_1.0 atlas (including N = 53 ICNs) (Du et al., 2020) to obtain the corresponding atlas in native space.

The ICNs of the normalized Neuromark_fMRI_1.0 atlas, which divides the brain into distinct regions, were used as nodes in the network to build the structural connectivity network. For each subject, we reconstructed an *N×N* structural connectivity network (M), counting the number of streamlines connecting pairs of nodes.

##### Evaluation criteria

First, to determine the effect of the defined objective function in our multimodal model, we used the same criteria adopted for the synthetic data. Consequently, for the ICNs of each model (sfCICA and CICA), the distance was estimated using **Eq. 9**. Then, we computed the sparsity of the estimated FNCs as in **Eq. 10**.

To enhance distance representation, we determined the difference of the measured distances, as shown in **Eq. 11.**

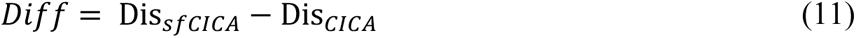

In addition, we evaluated the proposed model with real data, using a set of statistical analysis. A Paired t-test analysis was employed to determine if the examined ICNs from the sfCICA model, constrained by structural connectivity, differ significantly from those of the CICA model. The paired t-test analysis assesses whether the mean difference between paired observations is significantly different from zero. Here, we used the time course of the ICNs and their FNC matrices as observations for the paired t-test. FNC matrices were assessed using Pearson’s correlation among the time courses of the ICNs at the subject level for both the sfCICA and CICA models. Moreover, for group comparison between subjects with schizophrenia (SZ) and healthy control (HC), we applied a Generalized Linear Regression model (GLM) utilizing FNC matrices and spatial maps. GLM analysis is a statistical technique that provides a flexible framework for modeling the relationship between a dependent variable and one or more independent variables. In this study, we used GLM to regress out effects of the age, sex, diagnosis, motion, and imaging site effect as covariates for the FNC matrices and spatial maps. Additionally, false discovery rate (FDR) correction was performed to adjust the estimated p-values.

Finally, to show that structural connectivity helps tune the FNCs we used network graph parameters. To this end, graph theory analysis, utilizing FNC features was employed. We estimated the FNCs as Pearson’s correlation between TC related to each network (ICN) to construct the FNC network at the individual level for both the CICA and sfCICA. Then, we determined four common brain network parameters including modularity (Newman, 2006), local and global efficiency (Latora & Marchiori, 2001), and small worldness (Amaral et al., 2000) to characterize the graph properties of the resulting FNC. In addition, randomness (Vergara et al., 2018), and sparsity were also examined.

In a brain network, modularity indicates the presence of functional modules involved in specific cognitive processes (Newman, 2006). Local efficiency represents the efficiency of the communications between regions (here ICNs) and global efficiency is the capacity of the network for transferring information across the whole brain (Latora & Marchiori, 2001). Additionally, small-worldness is the ratio of clustering to shortest path length, which summarizes the balance between the efficiency and clustering in the brain network (Amaral et al., 2000).

## 3. Results

To determine multimodal ICNs through an iterative optimization, we applied the proposed sfCICA model to the diffusion MRI (dMRI) and the resting-state fMRI (rs-fMRI) data obtained from 30 synthetic samples and 311 real subjects (146 HC and 162 SZ). The number of the ICNs was determined based on a predefined standard ICN template, used as prior spatial maps. In this study, the NeuroMark_1.0 template was utilized as prior maps for the real data. This template consists of seven distinct functional domains, including subcortical (SC), sensorimotor (SM), auditory (AUD), visual (VS), cognitive control (CC), default mode (DM), and cerebellum (CB). Our results represent multimodal ICNs that are more informative and enhanced by incorporating multi-modal information. They provide supporting evidence indicating the impact of the structural connectivity information on neural activities (Litwińczuk et al., 2022).

### 3.1. Estimated ICNs using synthetic datasets show a structural and functional learning pattern

To determine the applicability and the robustness of the proposed model, we investigate the time course distances of the ICNs, as a measure of the correlation, for both the sfCICA and the CICA model at the subject level for different synthetic groups.

Distance analysis on the synthetic dataset predominantly revealed smaller distances after performing structural connectivity constraints using the sfCICA model compared to the CICA model, shown in **Figure 3**. This is reflected by average paired distances for the sfCICA (red box) and the CICA (green box) model. **Figure 3**, column **(a)** displays average paired distances at the subject level for each ICN. In all three groups of synthetic datasets, the average distances of the sfCICA were primarily constrained by structural connectivity and decreased compared to the CICA model. The corresponding group structural connectivity is shown in column (**b**).

**Figure 3.**
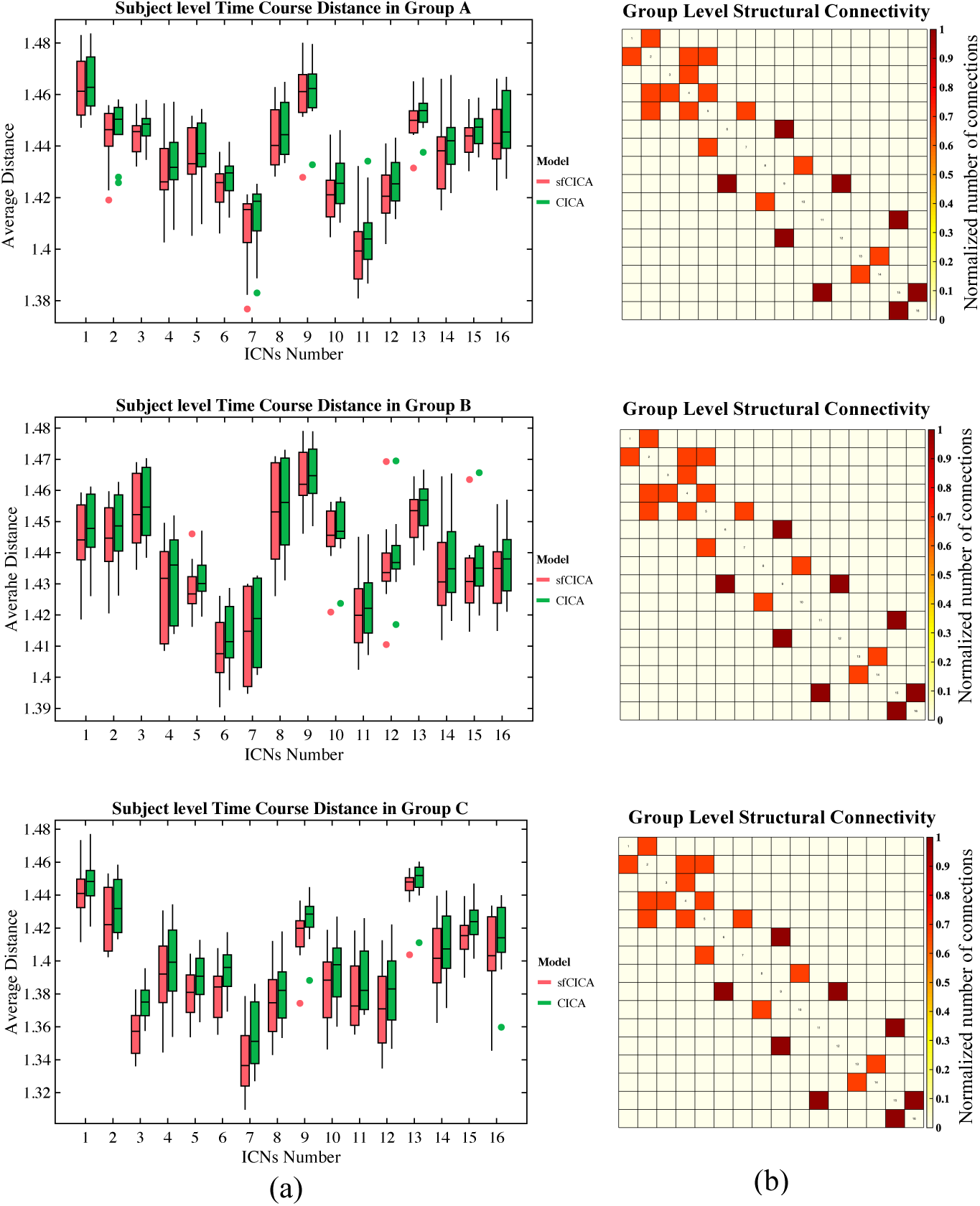
(a) Shows the average time course distance of the three groups of synthetic data, comparing the sfCICA (red box) to CICA (green box) at the subject level for each ICN. (b) Displays the corresponding group-level structural connectivity.

Statistical analysis, employing a paired t-test, revealed significant differences (p-value <0.05, FDR correction: q< 0.05) in the average group distances between the sfCICA and the CICA model for all the ICNs, except for ICN 14 in Group B, and ICN 1, 3, 7, 8, 11, 13, 14, 15 and, 16 in Group C.

Figure 4. represents the time course distance of an ICN (here ICNs 2 and 15) from other connected ICNs at the subject level across three different groups using both the sfCICA and CICA models. We specifically selected ICNs with varying connection weights: one exhibiting a high structural connection (ICN number 2) and another with a low structural connection (ICN number 15) to other ICNs. The results in Figure 4. illustrate a greater decrease in time course distance for the connected ICNs in the sfCICA model (red box) compared to the CICA (green box). As expected, in the subject-level analysis, the distance between ICNs in the sfCICA model decreased more significantly than those in the CICA model.

**Figure 4.**
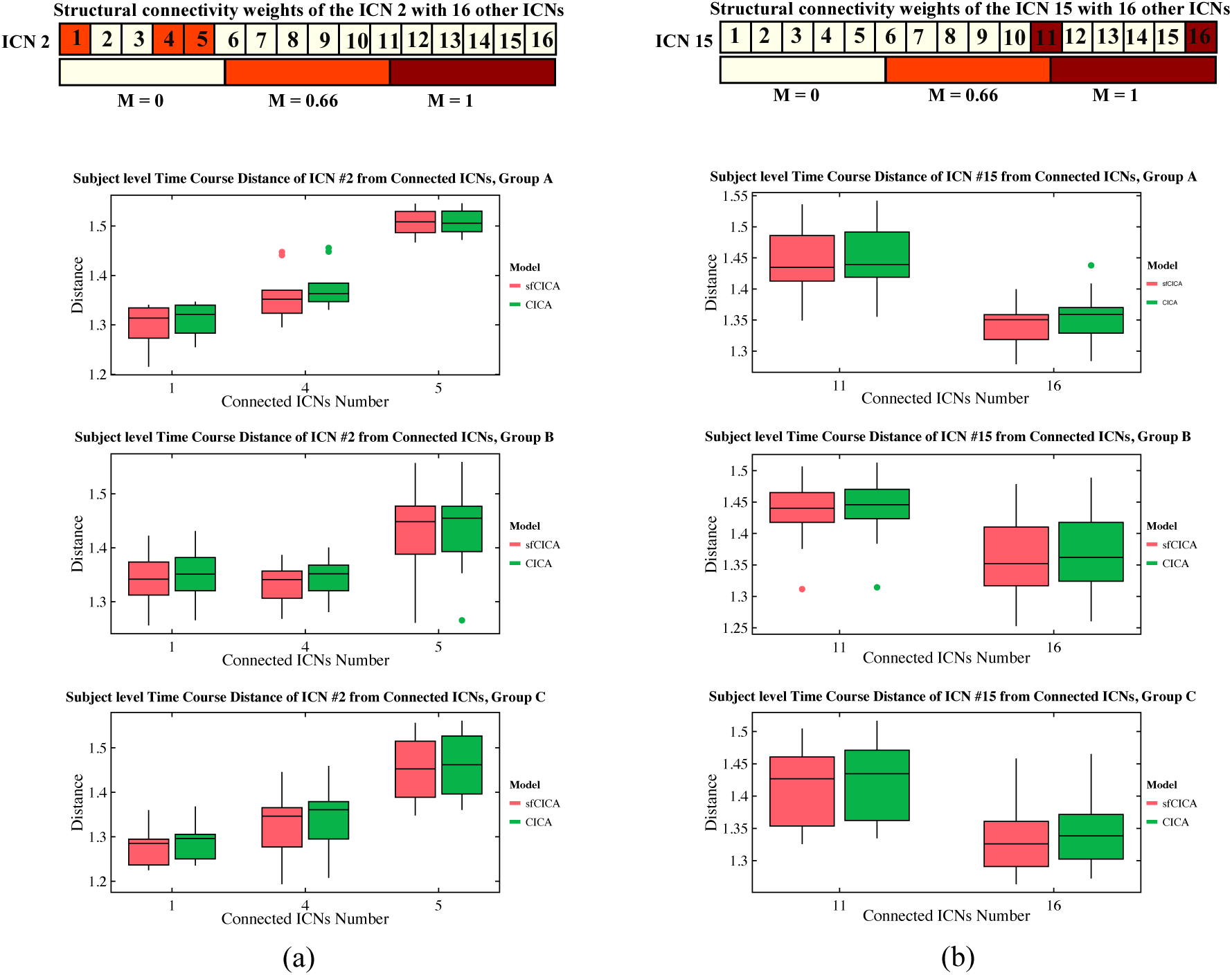
It depicts the estimated time course distance of ICN 2 and 15 with other connected ICNs at the subject level, shown in columns (a) and (b) respectively. The time course distance of the ICNs in sfCICA model was lower compared to CICA. The corresponding structural connectivity of ICN 2 and 15 with other ICNs comes at top of each column (a) and (b).

Moreover, we examined changes in distance specifically between two pairs of ICNs: one with a connection (between ICNs 2-4) and another without a connection (between ICNs 2-13), across three different synthetic datasets in Figure 5 (**a-c)**. The time course distance between ICN 2 - 4 (with strong structural connectivity, M = 0.66) and ICN 2-13 (without structural connectivity) was investigated at the subject level. Results show that the proposed model exhibits a lower distance compared to CICA for ICN 2-4 consistently across all subjects and various groups with different noise levels. However, for ICNs 2 - 13 (without structural connectivity), the time course distance for both sfCICA and CICA models was similar. The distance for the connected ICNs (e.g. 2-4) is higher than that for the unconnected ICNs (e.g. 2-13) when comparing the distance results between ICN 2-4 and ICN 2-13 at the subject level. The decrease in time course distance observed for connected ICNs in the sfCICA model is greater than in the CICA model. This reflects the influence of the structural connection and the adaptivity of our model in fluctuating the ICN estimation. For the non-connected ICNs (without structural connectivity), only functional information was effective, and the observed changes were consistently at a similar level for both models.

**Figure 5.**
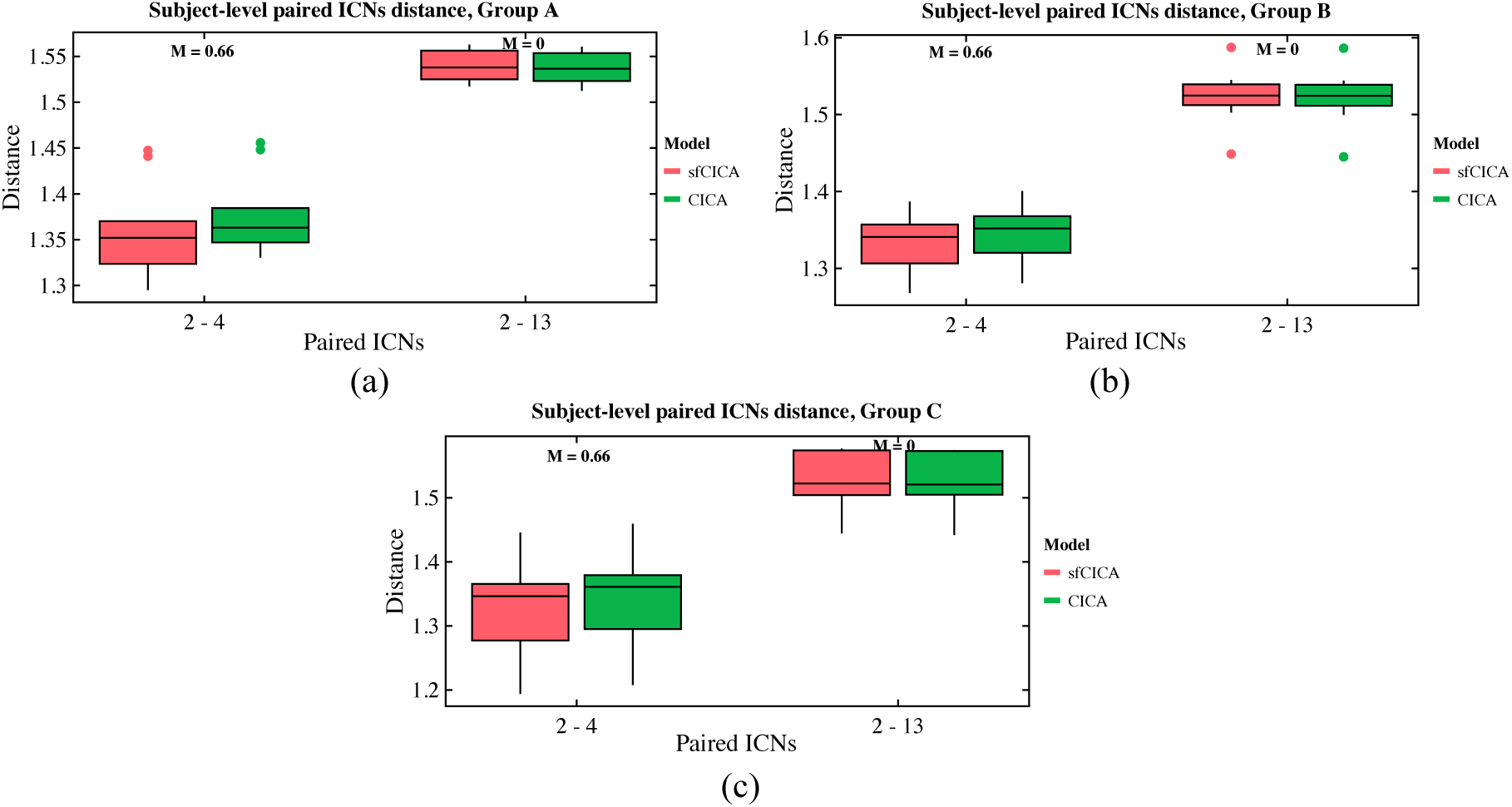
The time course distance of ICN 2 with a connected ICN (e.g 4) and a non-connected ICN (e.g. 13) is represented for synthetic data with three different noise levels **(a-c)**. The distance decreased for connected ICNs (e.g. ICN 2-4), while the disconnected ICNs had similar distances across all subjects.

Statistical analysis using a two-sample t-test reveals an increase in various graph metrics, (modularity: 0.352±0.01, global efficiency: 0.157±0.01, local efficiency: 0.199±0.009, small-worldness: 1.415±0.055, non-randomness: 5.0±1.15) for sfCICA compared to CICA (modularity: 0.351±0.01, global efficiency: 0.155±0.01, local efficiency: 0.198±0.009, small-worldness: 1.412±0.006, non-randomness: 4.18±1.15). A slight, but insignificant decrease (p-value>0.05) was observed between the sparsity measurements of the CICA (Mean±Std: 0.28±0.01) and sfCICA (Mean±Std: 0.27±0.01) models.

### 3.2. FNC is informed by structural connectivity weights in synthetic data

To emphasize the influence of the structural connectivity information on the FNC of ICNs in synthetic data, we assess the similarity of the FNC with structural connectivity, as well as the residual FNC for both the proposed sfCICA and the CICA models. The residual FNC is determined as the difference between the FNC matrices from the sfCICA and CICA models (FNC_diff_ = FNC_sfCICA_ - FNC_CICA_).

We perform two different evaluations by computing the similarity of the FNC matrix to determine the degree to which the structural connectivity impacted the resulting model weights, as shown in **Figure. 6**. For this purpose, the FNC matrix of the estimated ICNs from the sfCICA model was determined by computing Pearson’s correlation between paired ICNs for each subject. The similarity of the FNCs was then computed: (1) with structural connectivity and, (2) with FNC_diff_, at the subject level. A similar procedure was performed on the ICNs from the CICA model, and the results from both the sfCICA and CICA models were compared. From both evaluations, FNC_sfCICA_ demonstrated higher similarity with FNC_diff_ and structural connectivity than FNC_CICA_. These results suggest that the proposed sfCICA model is informed by structural information and remains robust to noise. Even with an increase in the noise level from Group A to C, the proposed model showed improvement compared to the CICA model. However, FNC_sfCICA_ and FNC_CICA_ exhibited the same level of similarity with structural connectivity.

**Figure 6.**
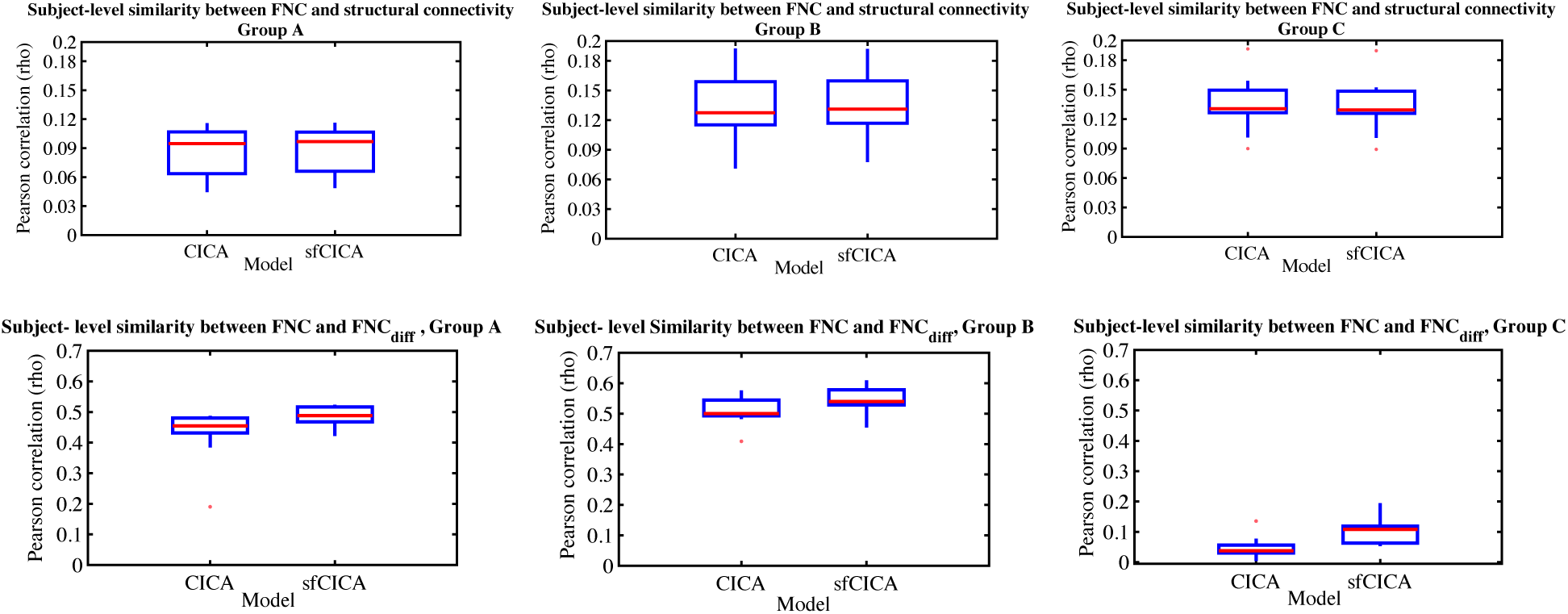
Similarity (Pearson correlation) of the FNC matrix with structural connectivity and with FNC_diff_ (FNC^diff^ = FNC_sfCICA_ - FNC_CICA_) was estimated for both the sfCICA and the CICA model across three distinct groups of synthetic data (columns). The results indicate that the sfCICA model exhibits a higher correlation with both structural connectivity and FNC_diff_ compared to CICA, as expected. Notably, greater differences were observed for FC_diff_. Furthermore, the findings demonstrate the robustness of the proposed model to noise, as evidenced by its consistent performance even as noise increases from Group A to C.

### 3.3. The subject-level structural-functional ICNs show more modular and integrated networks for the FBIRN data

Using the sfCICA model, the ICNs are guided by structural connectivity and FNC for each subject. In this paper, we used the NeuroMark template (Du et al., 2020) as prior spatial information, and the included information was evaluated by estimating the spatial similarity between the estimated ICNs and the corresponding NeuroMark template networks using Pearson’s correlation for both the sfCICA and the CICA models. To analyze the effectiveness of the proposed model for both healthy (HC) and disease (SZ) cases, we presented separate results for each group. In Figure 7a, the results showed an unimodal Gaussian distribution for HC and a bimodal Gaussian-like distribution in SZ, which is more clearly highlighted in the proposed model compared to the CICA model. It illustrates more significant spatial similarity for all subjects in the sfCICA model (Mean: 0.52) compared to the CICA model (Mean: 0.47) across both HC and SZ cohorts from the FBIRN dataset. On average, the sfCICA model exhibited a 10% increase in similarity compared to the spatial similarity estimated by the CICA model. The enhanced spatial similarity observed with standard and reproducible templates across multiple subjects suggests that informing the optimization process using structural connectivity could improve the results.

**Figure 7.**
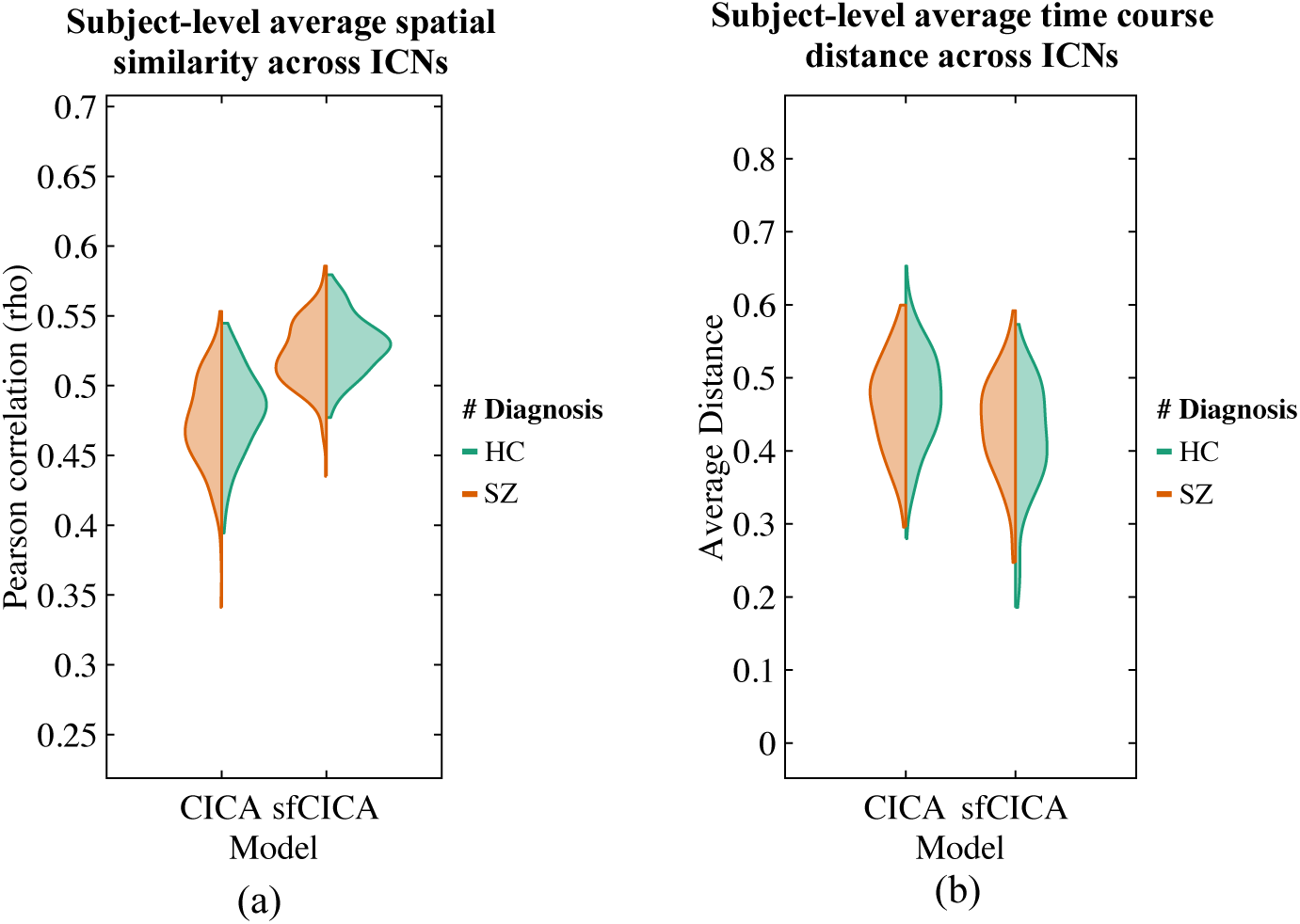
Distribution of average spatial similarity and time course distance over the ICNs at the subject level were represented for each model using FBIRN data. In (a), the distribution of the spatial similarity (Pearson correlation) with NeuroMark_01 template, used for prior spatial information, is illustrated for both the CICA and the sfCICA models in two groups: HC in green and SZ in red. Mean square distances of each time course from remaining time courses are shown in (b) within each HC (green) and SZ (red) group using the sfCICA and the CICA models. The sfCICA model showed higher spatial similarity and reduced time course distances.

Moreover, in Figure 7b, we computed and depicted the time course distance between paired ICNs at the subject level to assess the impact of structural connectivity. As we expected, the estimated distance showed a dependence on the structural connectivity weights, being consistently smaller for all the ICNs estimated by sfCICA in comparison to the CICA model. Overall, the distances, as well as the average distance of the time courses, were smaller in sfCICA than in CICA for both HC and SZ subjects, indicating greater functional integration (paired t-test: p-value<0.05). However, the increased spatial similarity resulted in more modular networks compared to the CICA model. In this context, distance refers to the mean square distances between paired time courses.

### 3.4. The subject-level structural-functional ICNs show a structural and functional learning pattern for the FBIRN data

Figure. 8 illustrates the differences in the estimated time course distances (a measure of the correlation) between the sfCICA and the CICA model for each ICN at the subject level. Negative distance values (below the pink line) indicate lower distances among ICNs in the sfCICA model compared to CICA. Interestingly, we observed predominantly negative correlations in the FBIRN dataset. In addition, ICN 18, 29, 37, 40, 4, 47, 48, 51, 52, and 53 show, on average, positive differences across all subjects. In general, we expect that ICNs with weak mean structural connectivity weight (as shown in **Figure. 8**) will show higher distances in the sfCICA model than in the CICA model, resulting in a positive average pattern for distance differences. **Figure. 8** reveals that the weights of the structural connectivity are associated with the distance of the time courses, i.e., where the structural connectivity is weak, the distance is large, and where the structural connectivity is strong, the distance is small among the sfCICA relative to the CICA time courses. These results suggest that both structural weights and functional spatial maps contribute complementary information to the estimated ICNs.

**Figure 8.**
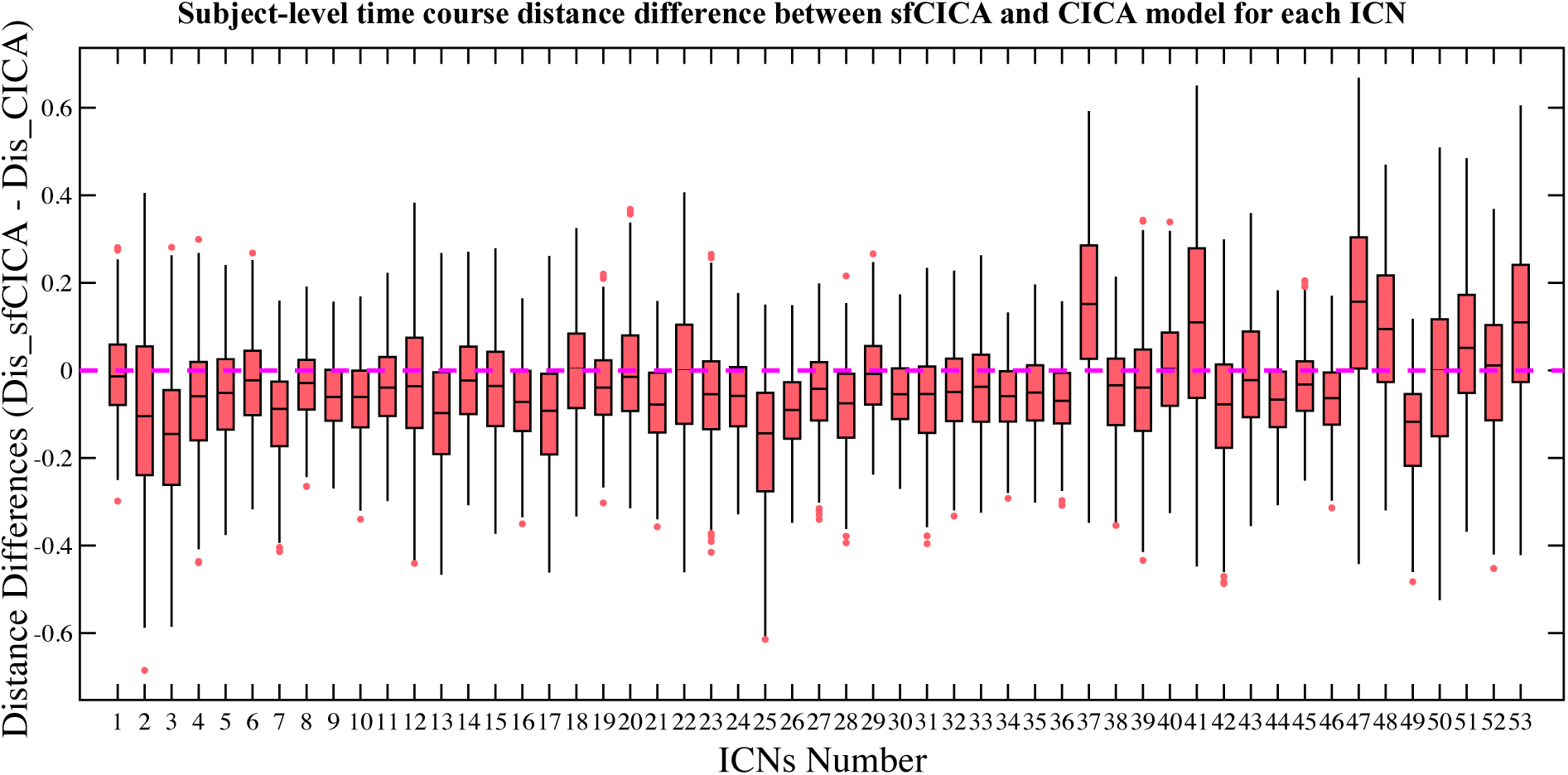
Illustrates that the differences in time course distances for most of the Intrinsic Connectivity Networks (ICNs) estimated by the sfCICA model, as compared to the CICA model in the FBIRN dataset, are smaller. The average distance differences of the time courses are predominantly negative, signifying smaller distances in the sfCICA compared to the CICA model, and this difference is influenced by Structural Connectivity.

We also estimated the time course distance for six randomly selected paired ICNs including 13-22, 48-47, 13-52, 12-48, 2-37, and 3-5, with different structural connectivity weights from weak to strong in the FBIRN data. As depicted in Figure 9, and similarly to the synthetic dataset, we observed distinct time course distances between sfCICA and CICA, with a decrease in sfCICA as the structural connectivity weights increased. On average over all subjects, our model (red box) has a higher distance for ICN 13 with 22 and 52 (both weak structural connections) compared to the CICA model (green box). For ICN 48, we consider two moderate (structural connectivity weight, M = 0.2) and weak connections (0.002) respectively with ICN 12 and 47. Interestingly, the distance between ICN 48 and 47 was increased compared to the CICA model, similar to the distance between ICN 13 and 52 (structural connectivity weight, M = 0.037). Conversely, the distance between ICN 48 and 12 was lower in our model. We evaluated the distance for the ICNs with strong structural connection, ICN 37 with 2 (M = 0.66) and ICN 3 with 5 (M = 0.91). As expected, for both connections, the distance decreased compared to the CICA model.

**Figure 9.**
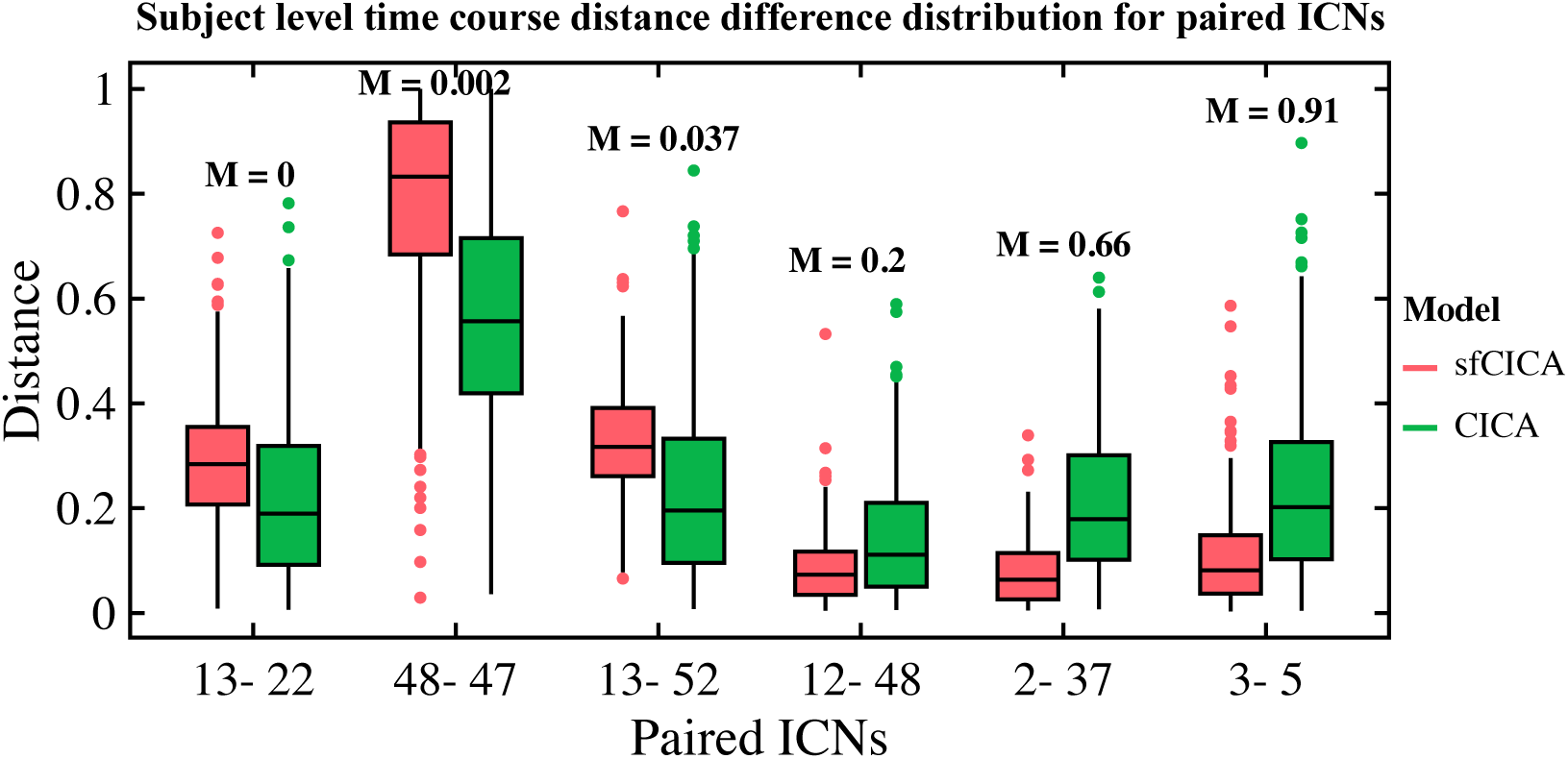
Depicts the distribution of time course distances for six paired ICNs estimated by both sfCICA and CICA models in the FBIRN dataset. These, paired ICNs were selected with varying structural connectivity weights ranging from weak to strong (0 - 0.91) connections. The estimated distance demonstrates an increase in structural connectivity weights, resulting in a decreased time course distance for the sfCICA model (red box) compared to the CICA model (green box). This highlights the influence of structural connectivity on the estimated ICNs.

### 3.5. Distinguished spatial patterns for the ICNs in the sfCICA compared to the CICA model using the FBIRN data

Significantly, we observed enhanced modularity and well-defined spatial maps in certain functional networks represented by ICNs IC 2, 22, 42, 7, 11, 21, 13,14, 34, 37, 50, 3, 27, 39, and 42, across most subjects, especially in those diagnosed with schizophrenia. These observations indicate that the structural connectivity constraints assist in better distinguishing between healthy and patient subjects within these specific ICNs. These functional networks mostly include subcortical, sensory networks (sensorimotor and, visual), default mode, and the cerebellum. In Figure 10, three different ICNs for subjects 29, 30, 37, and 74 are depicted in each row. All ICNs were converted to z-scores and thresholded at z-score>3, p-value<0.05. Additionally, a qualitative examination reveals more modular networks compared to the CICA model, (refer to Figure 11). Moreover, we performed paired t-test analysis between sfCICA and CICA using the spatial maps within each group (HC / SZ) separately. In healthy subjects, significant differences (p-value<0.05) were observed in most of the ICNs, except for the ICNs 45 and 51. ICNs 9, 27, and 51 in schizophrenia showed no significant differences between sfCICA and CICA.

**Figure 10.**
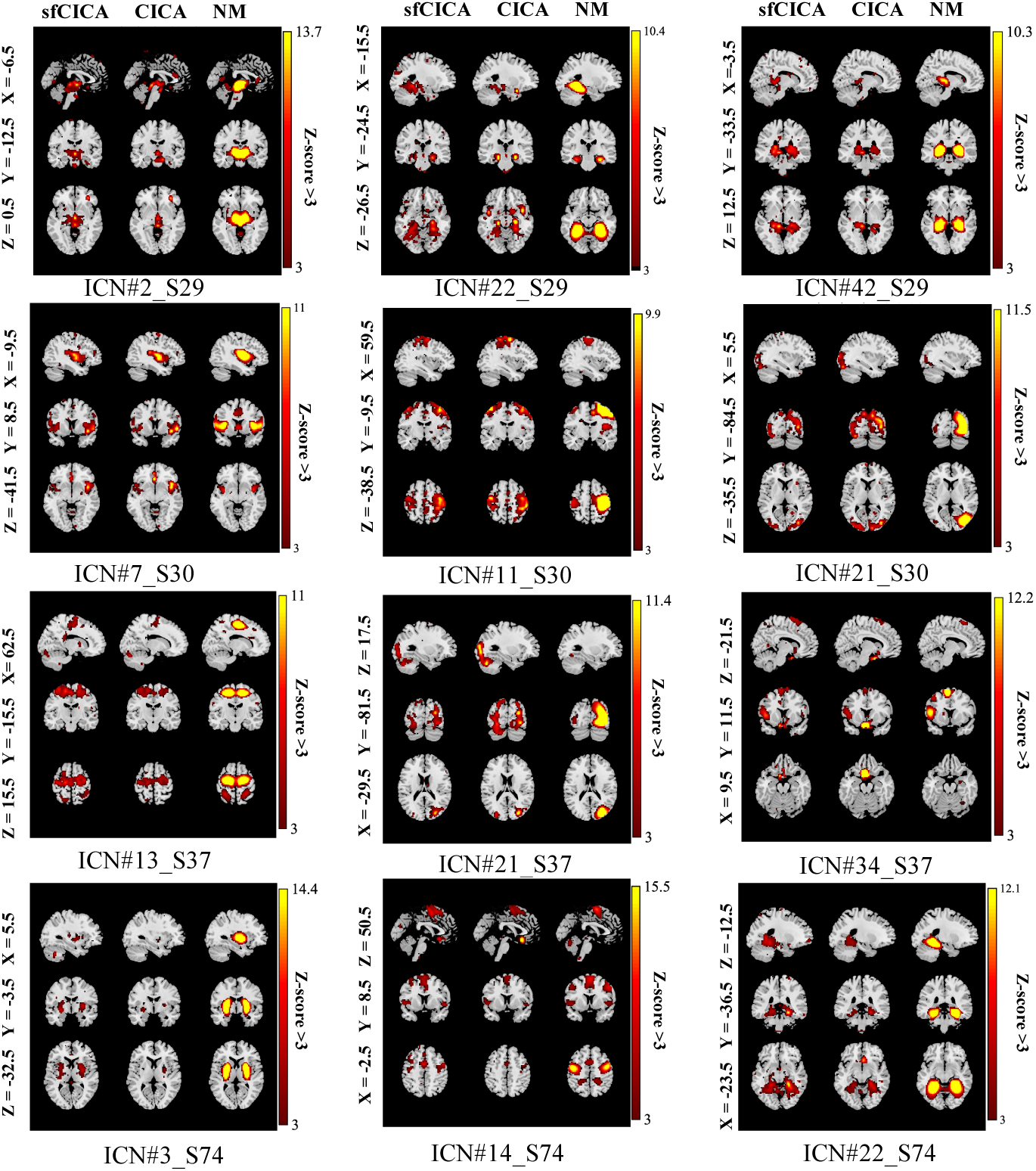
Each column displays spatial maps for selected ICNs (z-score>3) derived from different subjects using sfCICA, CICA, and the NeuroMark template (NM). The sfCICA consistently exhibits more modular characteristics (average modularity for sfCICA: 0.24, for the CICA: 0.21) across different subjects compared to the CICA model.

**Figure 11.**
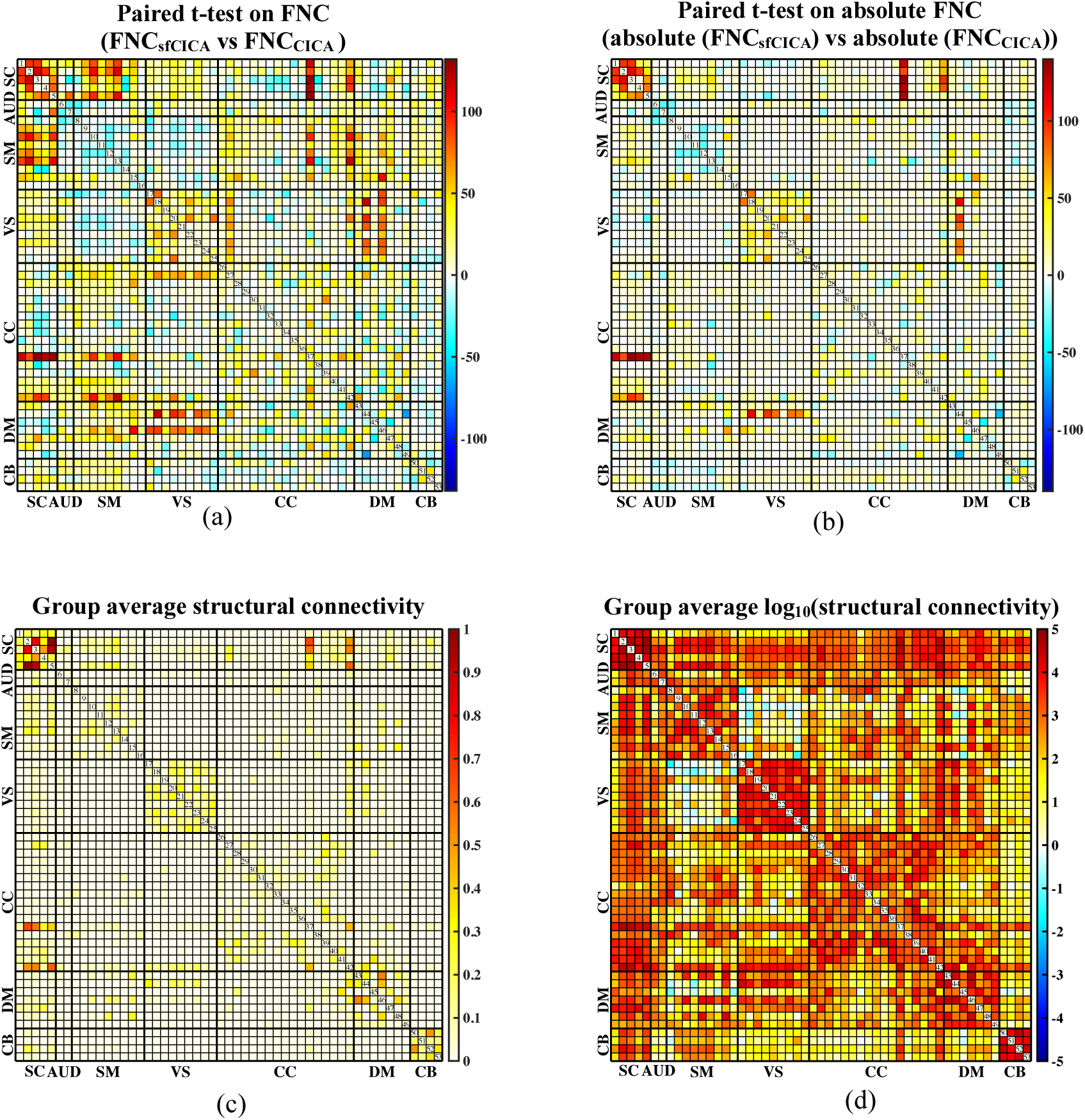
Illustrates the results for the paired t-test analysis between the FNC matrices of the sfCICA and CICA models, along with their corresponding structural connectivity, using FBIRN dataset. FNC represents the Pearson correlation between the estimated ICNs of each model. In (a). T-value maps for the paired t-test using FNC are shown, while (b) displays the t-map for the absolute FNCs. sfCICA model exhibits higher values in most ICNs compared to the CICA model. The group average structural connectivity, and its logarithmic scale connection weights (*log_10_(connection weights)*), are represented in (b) and (c). The weights of the structural connectivity indicate the number of tracts connecting two ICNs.

In addition, we computed graph-based network features such as modularity and randomness. Utilizing FNC matrices derived from the estimated Intrinsic Connectivity Networks (ICNs), our analysis revealed that the modularity coefficient for ICNs estimated with the sfCICA model was statistically significant (paired t-test: p-value < 0.05, Mean±std = 0.24±0.03) compared to the CICA model (Mean±std = 0.21±0.02). A randomness analysis was also conducted, indicating significant differences in non-randomness. Specifically, the sfCICA model exhibited a lower randomness coefficient (i.e., higher non-randomness characteristic) in the FNC patterns (paired t-test: p-value < 0.05, Mean±std = 13.3±3.5) compared to the CICA model (Mean±std = 11.5±2.05).

### 3.6. FNC of the Structural-functional connectivity informed ICNs revealed significantly constrained/ learned from structural connectivity using the FBIRN data

To determine the differences between the CICA and sfCICA models, we applied a paired t-test analysis on FNC matrices. Interestingly the paired t-tests indicated rejection of the null hypothesis, signifying significant differences between sfCICA and CICA across all ICNs. The estimated t-values (**Figure. 11a**) represented significant differences in ICNs related to the SC, SM, VS, CC, and DM networks. Paired t-test analysis was also performed regarding the absolute value for the FNCs, as illustrated in **Figure. 11b**, it shows reduced t-values for certain ICNs within the AUD, SM, VS, CC, DM, and CB networks. It replicates a decrease in the FNC association between the two models and can be explained based on the hypothesis that the impact of the relationship between structural connectivity and FNC on cognitive performance may depend on the functional domain.

Moreover, regarding **Figure. 11 (a-d)**, and the comparison of structural connectivity patterns (Figure 11**(c-d)**) with paired t-test analysis (Figure 11**(a-b)**), it is apparent that the structural connectivity weights optimize the distances, thereby enhancing the estimation of the ICNs. Further, analysis of the paired t-test between ICNs of both models reveals similarity for ICNs 1, 6, 11, 12, 14, 15, 18, 20, 22, 29, 39, 40, 43, 45, 50, and 52.

The determined results were jointly estimated by incorporating information from the structural connectivity weights into the correlation between the time courses overall time (i.e., the FNC matrix). A similar analysis on synthetic data was conducted on real data as well to assess the similarity of the FNC with structural connectivity and FNC with FNC_diff_, determining the extent to which structural connectivity impacts the resulting model weights, as illustrated in **Figure. 12**. For this purpose, an FNC matrix was computed for each subject using Pearson’s correlation among the time courses of the ICNs determined by the sfCICA model. A similar procedure was performed to estimate FNC_CICA_ using the ICNs of the CICA model. The comparison of results from both the sfCICA and CICA models indicated that in real datasets (FBIRN), FNC_sfCICA_ exhibits greater similarity with structural connectivity and FNC_diff_ to FNC_CICA_. It indicates the proposed sfCICA model is informed by structural information, showing the multi-modal sfCICA model provides additional information that could capture variability between subjects and between groups.

**Figure 12.**
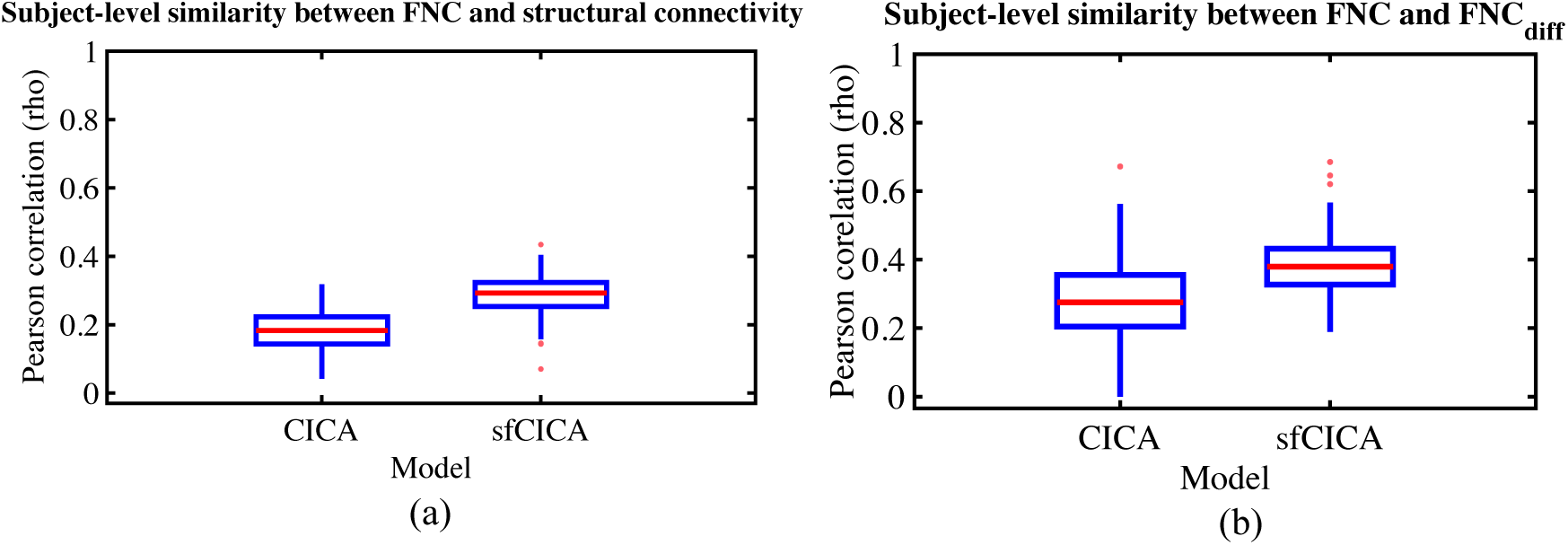
Similarity (Pearson correlation) between FNC matrix of estimated ICNs from the sfCICA model with structural connectivity and, residual (diff) FNC matrix is represented for FBIRN dataset respectively in (a) and (b). Results show similarity (correlation) with both structural connectivity and residual FNC (FNC_diff_) is higher for the sfCICA model than the CICA, representing more informatic ICNs.

### 3.7. Structural-functional connectivity-informed ICNs reveal significant differences between controls and schizophrenia subjects using the FBIRN data

Figure. 13 (**a**-**b**) show the average FNC across all subjects for the sfCICA and CICA models. Additionally, group differences analysis using FNC for the HC vs SZ was performed and the results are represented in **Figure. 13 (c-d).**

**Figure 13.**
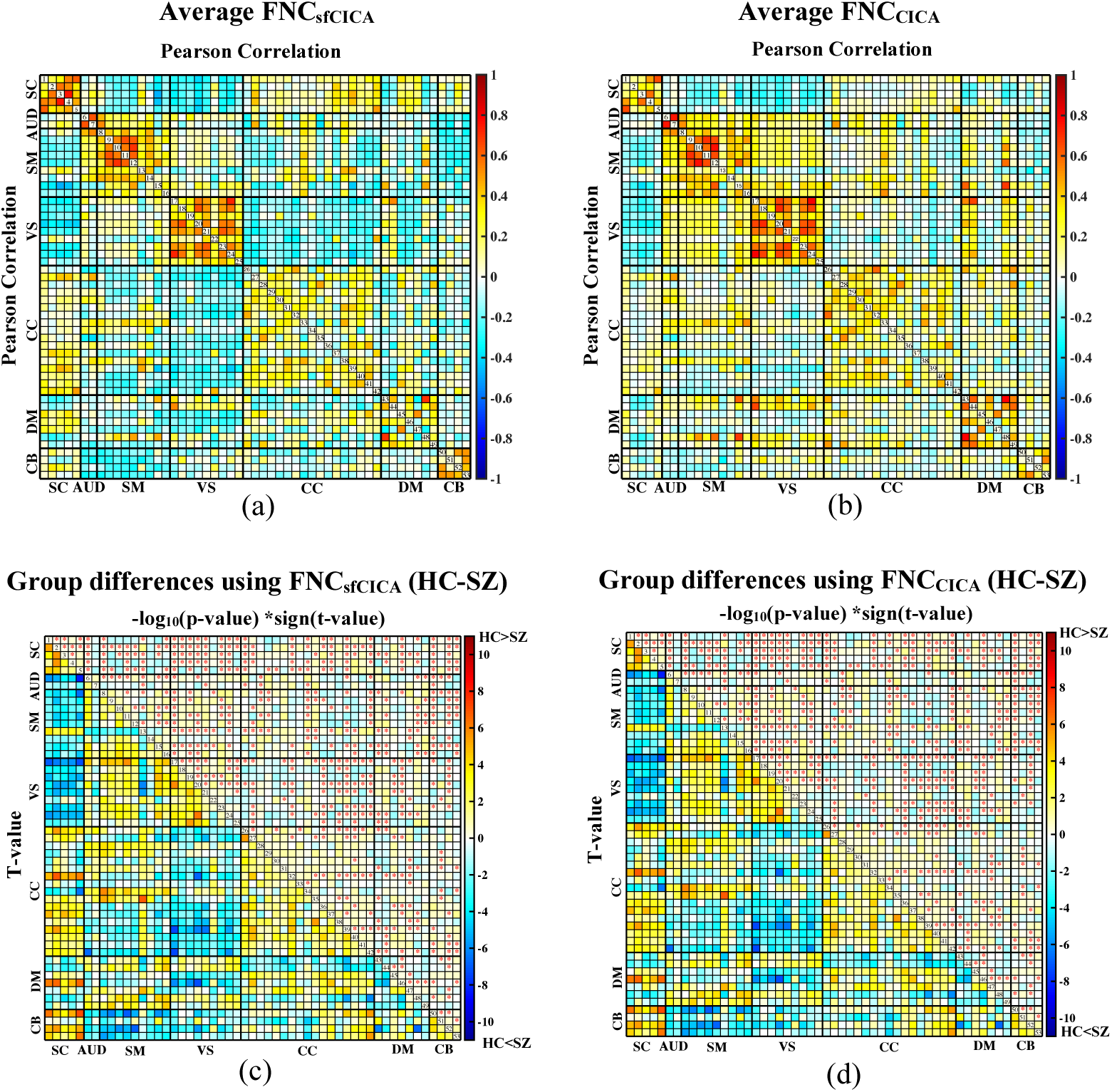
Average FNC estimated across all subjects and statistical group difference maps (HC/ SZ) for the FBIRN dataset. In **(a)** and **(b)**, the average FNC of all subjects is presented for the sfCICA and CICA models respectively. For the group (HC/ SZ) difference comparison, GLM analysis was performed to regress out covariates including age, sex, motion, imaging site, and diagnosis from the FNC matrices. More significant differences (p-value<0.05 and FDR<0.03), represented by red stars, were observed between HC and SZ in the proposed sfCICA compared to the CICA model. Results were corrected for multiple comparisons using false discovery rate (FDR) with q<0.03; and represented in **(c-d)**, which upper triangle each cell of the matrices are *-log_10_(p-value) *sign(t-value)* and lower triangle is t-value. ICNs were categorized in seven different domains including subcortical network (SC), auditory network (AUD), sensorimotor network (SM), visual network (VS), cognitive control network (CC), default mode network (DM) and cerebellum (CB). Results show that FNC in HC are higher than SZ in most of the ICNs, in the sfCICA compared to the CICA model.

Furthermore, to investigate the ability of the proposed method to find group differences (healthy/ patient) in FNC patterns using both the sfCICA and the CICA models, a generalized linear regression model (GLM) was performed with multiple covariates (**Figure. 13** (**c**-**d**)). The results indicated a more significant correlation (p-value<0.05, FDR: q<0.03) across all networks (displayed with red stars) comparing HC vs SZ group for the sfCICA model. We detected 503 significant cells in the upper triangle for the sfCICA model (Figure 13 **c**), a notable increase compared to the 421 significant cells observed for the CICA model (Figure 13 **d**). Correlations of the sensory networks were less significant in the sfCICA model compared to the CICA. In contrast, SC-related regions showed more significance. The findings suggested the involvement of structural connectivity in the SC domain, a factor not observed with the CICA model. ICNs for the SC, DMN, and CB networks exhibited significantly higher FNC in HC compared to SZ for the sfCICA model (**Figure. 13 c**), while sensory-related networks (AUD, VS, and SM) and CC network showed significantly higher FNC in HC compared to SZ for both models (**Figure. 13 (c-d)**). This suggests that considering structural connectivity plays an important role in identifying the FNC of the SC and that the sfCICA which jointly analyzes structural and functional ICA output provides a more sensitive model. The values illustrated in the higher triangle of matrices in Figure 13 **(c-d)**, are −log10(*p*_*value*) × *sign* (*t*_*value*) and t-values are represented in the lower triangle.

### 3.8. Multimodal ICNs represent more significant spatial maps between groups (controls and schizophrenia) using the FBIRN data

To investigate the hypothesis of the proposed model, we conducted a group differences analysis between HC and SZ on the ICN spatial maps. For this purpose, a GLM analysis was adopted to regress out the effect of age, sex, motion, site, and diagnosis per ICN. Table 1 displays the percentage of significant voxels in each ICN of different networks. Results indicate that out of fifty-three ICNs, thirty-two showed a higher number of significant voxels (p-value<0.05) in the sfCICA model (dark blue rows). Conversely, in the remaining twenty-one ICNs, five (light blue row) demonstrated a similar percentage of significant voxels, and sixteens (white rows) represented a lower percentage with a deviation for the sfCICA model.

**Table 1.** Shows the percentage of the significant voxels in each ICN using group differences analysis. ICNs (rows) with higher values for the sfCICA are represented in dark blue, while light blue and white indicate equal or lower values for the sfCICA compared to the CICA.

| Networks | ICNs | Number of significant voxels using CICA (%) | Number of significant voxels using sfCICA (%) | Differences between significant voxels, sfCICA – CICA (%) |
| --- | --- | --- | --- | --- |
| SC | IC1 | 4.9 | 5.6 | 0.7 |
|  | IC2 | 4.1 | 6 | 1.9 |
|  | IC3 | 4.2 | 4.9 | 0.7 |
|  | IC4 | 3 | 3.5 | 0.5 |
|  | IC5 | 8.3 | 9.6 | 1.3 |
| AUD | IC1 | 2.8 | 2.9 | 0.1 |
|  | IC2 | 4.7 | 5.4 | 0.7 |
| SM | IC1 | 2.1 | 2.9 | 0.7 |
|  | IC2 | 2.8 | 3.4 | 0.6 |
|  | IC3 | 7.1 | 5.7 | -1.4 |
|  | IC4 | 8.3 | 9.1 | 0.8 |
|  | IC5 | 5.5 | 5.8 | 0.3 |
|  | IC6 | 4.4 | 4.2 | -0.2 |
|  | IC7 | 3.9 | 3.9 | 0 |
|  | IC8 | 6.6 | 7 | 0.4 |
|  | IC9 | 5 | 5.1 | 0.1 |
| VS | IC1 | 3.2 | 4.1 | 0.9 |
|  | IC2 | 10 | 10 | 0 |
|  | IC3 | 5.1 | 5.2 | 0.1 |
|  | IC4 | 8 | 6.8 | -1.2 |
|  | IC5 | 3.5 | 4.2 | 0.7 |
|  | IC6 | 3.1 | 3.4 | 0.3 |
|  | IC7 | 3.8 | 4.5 | 0.7 |
|  | IC8 | 2.4 | 2.4 | 0 |
|  | IC9 | 4.1 | 4.8 | 0.7 |
| CC | IC1 | 5.1 | 5.1 | 0 |
|  | IC2 | 8.2 | 7.3 | -0.9 |
|  | IC3 | 5.5 | 5.9 | 0.4 |
|  | IC4 | 5.3 | 5.2 | -0.1 |
|  | IC5 | 6.6 | 6.7 | 0.1 |
|  | IC6 | 6.7 | 6.2 | -0.5 |
|  | IC7 | 5.2 | 6 | 0.8 |
|  | IC8 | 8.9 | 6.3 | -2.6 |
|  | IC9 | 8 | 8.3 | 0.3 |
|  | IC10 | 7 | 6.4 | -0.6 |
|  | IC11 | 9.2 | 8 | -0.8 |
|  | IC12 | 3.3 | 3.7 | 0.4 |
|  | IC13 | 8 | 8.3 | 0.3 |
|  | IC14 | 4.3 | 5.5 | 1.2 |
|  | IC15 | 7 | 5.4 | -1.6 |
|  | IC16 | 4.2 | 4.2 | 0 |
|  | IC17 | 5.6 | 5.3 | -0.3 |
| DM | IC1 | 3.1 | 3.4 | 0.3 |
|  | IC2 | 3.5 | 3.2 | -0.3 |
|  | IC3 | 2.5 | 3.2 | 0.7 |
|  | IC4 | 8.2 | 8.1 | -0.1 |
|  | IC5 | 4.5 | 3.6 | -0.9 |
|  | IC6 | 5.1 | 5.2 | 0.1 |
|  | IC7 | 7.6 | 6 | -1.6 |
| CB | IC1 | 2.1 | 4.8 | 2.7 |
|  | IC2 | 9.1 | 6.4 | -2.7 |
|  | IC3 | 2.1 | 5.9 | 3.8 |
|  | IC4 | 4.6 | 5.6 | 1 |

In addition, network properties including modularity, efficiency, small-worldness, randomness, and sparsity were estimated using FNC features. Results (Table 2) showed significantly higher (p-value<0.05) efficiency, modularity, and randomness for the sfCICA model using the FNC feature. Small-worldness was also greater for sfCICA but the difference between the two models was not significant.

**Table 2.** Graph parameters estimated for the FNC matrices.

| Graph parameters | Single-modality Model<br>(Mean±std) | Multi-modality Model<br>(Mean±std) |
| --- | --- | --- |
| Modularity | **0.21±0.02** | **0.24±0.03** |
| Local efficiency | **0.32±0.01** | **0.37±0.01** |
| Global efficiency | **0.30±0.03** | **0.33±0.03** |
| Small-worldness | **1.15±0.08** | **1.17±0.06** |
| Non- Randomness | **11.30±3.50** | **13.50±2.05** |
| Sparsity | **0.28±0.01** | **0.27±0.01** |

## 4. Discussion

This study proposes a new, joint structural-functional, data-driven model to estimate intrinsic functional connectivity networks (ICNs) using multi-modal brain images. While most of the studies typically rely on fixed regions weighted by diffusion MRI (dMRI), our proposed approach is a multi-modal ICA model, named sfCICA, which is guided by the structural and functional connectivity network domain. Using network science, structural and functional information is compressed in nodes and edges which has become a common practice in neuroscience to understand functional interactions in the brain (Babaeeghazvini et al., 2021). In the sfCICA model, the functional connectivity from resting-state magnetic resonance imaging is constrained by structural connectivity weights (normalized number of streams). We applied the proposed model to datasets from subjects with schizophrenia and healthy controls, as well as simulated data. Through this new model, we identified improved spatial maps, individual subject variability, and modularity of the FNC networks. Furthermore, the sfCICA model exhibits less randomness and greater sensitivity to group differences in healthy controls and subjects with schizophrenia (HC/SZ subjects).

Most previous ICA models are focused on functional information or functional information with prior spatial constraints (Du & Fan, 2013; Du et al., 2020). In recent years, there has been a shift towards multi-modal models, incorporating features estimated from different modalities through joint analysis (Wu & Calhoun, 2023). Joint-ICA, parallel ICA, link-ICA, and cmICA models utilize features from fMRI, dMRI, or sMRI, such as connectivity, fractional anisotropy, and structural measures to identify functional networks. The proposed model in this study diverges by simultaneously learning from both functional and structural connectivity. This approach aims to constrain functional changes over time based on estimates of structural connectivity.

Less work has been given to dMRI connectivity, as in the cmICA model (Wu & Calhoun, 2023). cmICA (Wu & Calhoun, 2023), uses multi-modal fMRI and dMRI, integrating features derived from each modality to identify ICNs shared between structural connectivity and FNC, acknowledging that there might be a mismatch between features from each modality. Although cmICA attempts to address this, the contributions of features from each modality are asymmetrical (Wu & Calhoun, 2023). In contrast, our proposed model is based on both direct and indirect features and the contribution of each modality enhances the other. Thus, when structural connectivity weights are strong, functional data exhibits a higher correlation, while weaker structural connectivity weights result in less correlated functional data. Notably, the sfCICA model relies on two main concepts of an ICA analysis, maximizing independence and similarity, and tries to estimate intrinsic brain functional networks which are optimized based on both structural and functional connectivity information. Therefore, we posit that the proposed joint multi-modal information integration allows for better characterization of healthy and disordered brain connectivity (Iraji et al., 2016; Wu & Calhoun, 2023). Characterization of the intrinsic brain activity, known as functional network modeling, has been widely used in human brain studies, particularly because of its relevance to brain disorder research (Wang et al., 2021). One of the greatest advancements in this area was uncovered by rs-fMRI images using ICA analysis (Calhoun et al., 2001) and continues to advance for the field by the inclusion of additional imaging modalities.

Including two different modalities of information in our model, the replicability and noise robustness of the proposed model were assessed through three different synthetic datasets (**Figures 3, 4, and 5**). Each group has different levels of signal-to-noise ratio (SNR), and for each group, a similar pattern, constrained by structural connectivity, was observed. Our findings highlighted the effect of the structural connectivity constraints on the time courses, as their distances were significantly smaller when structural connectivity was stronger. Additionally, by changing the functional information and keeping the structural connectivity constant, the learning procedure is dependent on information from both modalities during the estimation of ICNs.

Empirical results from our proposed multi-modal model on real data show that our model (the sfCICA) is more sensitive in detecting functional connectivity in both healthy and schizophrenia groups, compared to the CICA model. Estimated ICNs were significantly more spatially similar (Figure 7a) to the reproducible template (NeuroMark template (Du et al., 2020)) for both healthy and schizophrenia subjects. This may enhance the replicability of the components using different datasets (Duda et al., 2023).

Moreover, the proposed sfCICA model aims to estimate a unified and integrated brain network using regional functional information instead of FNC as in (Zhu et al., 2021) which is the first time this has been performed in an ICA analysis. Overall, the results in Figure 7b showed that the distance of the time courses was smaller, which can be interpreted as more functionally correlated, and hence more uniform ICNs. While considering each ICN specifically (**Figures 8 and 9**), ICNs related to cognitive control (CC), and cerebellum (CB) networks were less constrained. In other words, distance minimization of them was less compared to other ICNs. It is highlighted that overall, they are less structurally connected to other ICNs. For instance, ICNs number 37 and 41 in the CC network, represent the weakest connection with CC, SM, and CB networks. Also, consistent with prior findings (Glasser et al., 2016; Van Essen et al., 2019; Wu & Calhoun, 2023), the DM has the largest distances within its network ICNs and the weakest with intra-network such as SC network. These findings suggest the advantages of multi-modal approaches, using a combination of structural and functional features, in facilitating the interpretability of the results which is consistent with previous findings (Glasser et al., 2016; Van Essen et al., 2019).

Alternatively, there is a hypothesis that structural connectivity may directly influence FNC, as higher structural connectivity leads to higher FNC (Litwińczuk et al., 2022; Zhao et al., 2023). Statistical analysis on FNC of the estimated multi-modal (sfCICA) ICNs and single model ICNs (CICA) in **Figure 11 (a-b)**, represented no association between the FNCs of the two models in specific domains including SM, AUD, VS, CC, and DM. Interestingly results were following previous findings (Li et al., 2020; Tsai et al., 2019). There is evidence that structural connectivity and FNC have no coupling in SM and VS, which means either structural connectivity is not dense, or it may come from dysfunction characteristics of the SZ subjects accompanied by weaker structural connectivity in SM, anterior cingulum cortex (ACC, related to DM domain) (Li et al., 2020). So, our results revealed it in the estimated ICNs features (FNC) using the sfCICA model. Regarding the VS domain, little is known about it, however, it has been shown that the connectivity of the visual system in SZ would be reduced (Reavis et al., 2020) which is consistent with our findings. Overall, findings suggested that integrating multi-modal information to estimate ICNs could help to find more accurate intrinsic functional domains in comparison to single-modality models such as the CICA.

Our proposed model (sfCICA) was more sensitive to detecting differences between individuals with schizophrenia patients and healthy controls. In schizophrenia both structural and functional networks are affected (Li et al., 2017; S. Qi et al., 2019), our results showed a broad range of impacted networks. Regarding the FNC matrices (Figure 13 a-b), findings are consistent with, and extend, those of prior studies. For example, prior studies have reported dysfunction in the CC network, which was also observed in our statistical results using the single modality CICA model. However, the proposed sfCICA model showed enhanced functional connectivity within the SC network with most of the domains, the CC network with other domains (e.g., visual), and the DM network as well as with SM, AUD, and VS. Overall, these findings suggest that the contribution of structural connectivity information could enhance our ability to identify dysfunctional connections. Also, ICNs related to memory and higher-order cognitive control functions related to DM networks have been reported as affected regions in individuals with SZ (Aleman et al., 1999; Minzenberg et al., 2009; Potkin et al., 2009; S. Qi et al., 2019). Consistent with prior findings our results show that functional correlations of the DM network are lower for the positive connections and higher for the negative connections (S. Qi et al., 2019), representing the effectiveness of our multi-modal model. However, in the proposed model, mostly FNC is higher for healthy compared to schizophrenia but there exist some ICNs that do not show the same pattern. For example, in ICNs 8 (of the SM network) and 42 (of the CC network), FNC is higher for schizophrenia. Compared to previous studies, mostly dysconnectivity occurs in DM and SM networks (Rong et al., 2023), and FNC increment is observed in these domains.

There exist some limitations in this study which would be interesting to consider in future works. We selected the model order using a standard ICN template including 53 ICNs (NeuroMark_1.0). The identified ICNs are restricted to prior spatial maps and are modified spatially and temporally during the optimization process. However, it might be variable over subjects, which should be considered in future works. Moreover, the standard template is functionally defined. Our multi-modal sfCICA model uncovers differences in functional connectivity between regions and their interactions are affected by structural information. To include differences in spatial maps and enable the comparability of results, a multi-modal (structural/functional) template would be needed. In parallel, increasing structural and functional studies highlights the need for a multi-modal simulator. Our synthetic data are independent and structural information has no impact on fMRI simulation. It could be beneficial to develop a simulator for generating structurally informed fMRI data (e.g. perhaps via a copula framework (Silva et al., 2014)). In addition, our current model starts from fixed networks for the structural connectivity, it would be useful to extend the model to also allow for adaptive updates to the structural connectivity at the single subject level, similar to what is occurring with the fMRI data. Existing methods have used structural connectivity as a deterministic constraint to functional connectivity (Seguin et al., 2022). In contrast, in our work, we propose a data-driven multi-objective model using structural connectivity constraints to incorporate these into an ICA model resulting in spatial maps, time courses, and FNC. Our expectation is that the results would increase our sensitivity to patient vs control differences. Our suggested model is one of the first tools for directly linking structural connectivity/FNC in a data-driven analysis. Furthermore, future extensions can be developed that allow the structural connectivity model to adapt to the functional data in the context of a supervised model, optimizing structural connectivity/FNC to maximize group differences and then using this in a new dataset to classify. Other future work could be to use our model with an alternative transformation of the structural connectivity network. Other future work could be to use our model with an alternative transformation of the structural connectivity network. Most of the not directly connected regions have polysynaptic communication (Seguin et al., 2022). One way to guide polysynaptic signals in structural connectivity/FNC models is by using communication network models (Seguin et al., 2022). Communication network models are derived from structural connectivity and converse to the structural connectivity, communication network models are dense (Seguin et al., 2022). One limitation of data fusion is that the structural connectivity informed FNC may be more complex to interpret than the FNC estimated from fMRI alone. However, there are also advantages as we point out, especially in enhancing sensitivity to group differences. One way to address this is to perform both analyses as we have done. An advantage of the use of the NeuroMark spatially constrained ICA framework is that the data-driven networks maintain their correspondence with one another due to the similar spatial priors in both. On the other hand, regions-based structural connectivity was identified before optimization, and it remains constant during resting-state data acquisition. In addition, for future works, it might be more informatic to use diffusion spectrum imaging (DSI). Although time courses and spatial maps are updated, the model can be extended to modify structural weights throughout the optimization process. Beyond this, we can also incorporate the dynamics of brain function.

## 5. Conclusion

In this work, we proposed a novel, structural-functional constrained independent component analysis (sfCICA), which incorporates both structural and functional connectivity information, guided by spatial maps to estimate functional networks in the human brain. The main motivation for the proposed model is to allow for structural and functional information to jointly influence the model estimation. The resulting ICNs were more spatially focal, and more synchronized in the sfCICA model compared to the CICA model. In addition, we observed that integrating structural connectivity information in the sfCICA enhances the sensitivity to group differences. The results were consistent with those based on synthetic structural and functional imaging data, with different functional information and noise levels. In sum, our findings demonstrate the effectiveness of the proposed model in both synthetic and real data. The approach applies to the study of a wide range of areas including brain development, aging, healthy control, and mental disorders.

## 6. Acknowledgment

This work is funded in part by the NSF grant 2112455 and the NIH grants R01MH118695 and R01MH123610.

## 7. Authors contribution

Mahshid Fouladivanda: Conceptualization; Formal analysis; Investigation; Methodology; Software; Visualization; Writing – original draft; Writing – review & editing. Armin Iraji: Conceptualization; Formal analysis; Writing – review & editing. Lei Wu: Software. Theodorus G.M. van Erp: Resources; Writing – review & editing. Aysenil Belger: Resources. Faris Hawamdeh: Writing – review & editing. Godfrey Pearlson: Writing – review & editing. Vince Calhoun: Conceptualization; Funding acquisition; Investigation; Methodology; Project administration; Resources; Software; Writing – review & editing.

